# The T-cell receptor repertoire of wild mice

**DOI:** 10.1101/2025.08.08.669451

**Authors:** Jacob Cohen, Simon Hunter-Barnett, Gayathri Nageswaran, Suzanne Byrne, Gemma Freeman, Matthew Cowley, Benny Chain, Mark Viney

**Affiliations:** Department of Evolution, Ecology and Behaviour, University of Liverpool, Liverpool, L69 7ZB, UK; Division of Infection and Immunity, University College London, WC1E 6BT, UK

## Abstract

Wild animals live in a pathogen-rich environment, and are normally infected with a wide range of micro- and macro-parasites. Wild animals’ T cells are central to the effectiveness of their adaptive immune response in ameliorating the effect of these infections. Here we have investigated the T-cell receptor (TCR) repertoire of wild mice to investigate how it varies in animals of different ages and sex, and from different sites. We sequenced the TCR alpha and beta chains of CD4^+^ and CD8^+^ T-cells of 65 wild *Mus musculus domesticus* from two UK sites. We analysed repertoire richness and diversity finding that wild mice have large TCR repertoires. Repertoire richness, which measures the breadth of the repertoire, was not significantly affected by mouse age or sex, suggesting that wild mice maintain the capacity to respond to novel antigens throughout their lives. In contrast, repertoire diversity (measured by Shannon’s index) was affected by a mouse sex-by-age interaction. This low diversity, coupled with constant richness, points to older mice having comparatively more highly abundant clones in their repertoires, perhaps due to chronic exposure to persistent pathogens in their environment. These findings provide a novel description of the wild mouse TCR, revealing an immune system that balances maintaining a broad response capacity with developing strong, lasting responses to infections in the natural environment.

## INTRODUCTION

The vertebrate immune system protects individuals from infection and disease. Most of our description and understanding of this system comes from study of the laboratory mouse, but laboratory mice aren’t exposed to the infections and other threats of the natural environment, which is the context in which the immune system and its function have evolved. There has been recent interest in studying the immune system and immune state of wild mice and other rodents (Viney & Riley 2017), finding that in wild mice the immune system is in an altered, often more activated, state compared with that of laboratory animals (Beura *et al*. 2016; Abolins *et al*. 2017; Abolins *et al*. 2018). Other studies have used rewilded mice, that is laboratory mice released into outdoor enclosures, and have found that they have an altered immune response that, for example, makes them susceptible to a parasitic nematode infection, while laboratory-maintained mice are resistant to it (Leung *et al*. 2018). Transplanting laboratory mice with wild mouse microbiomes also brings about profound immunological changes, reducing inflammation and, for example, increasing susceptibility to infection with influenza virus (Rosshart *et al*. 2017). The unifying thread from these studies is that laboratory mice are underexposed to infections compared with wild / rewilded / wild microbiome-exposed mice, and that greater infection stimulates the immune system into a different functional state, compared with laboratory mice.

B and T-cells are the principal players of the adaptive immune response and the dominant cell types in the lymphatic system, together (in humans) accounting for 40 % of immune cells (*c.* 25 % T-cells, 15 % B-cells) (Sender *et al*. 2023). T-cells bind to antigenic peptide molecules that cause the T-cells to proliferate and then to act directly against the antigenic source. Antigens are perceived by heterodimeric (in mice and humans principally α and β chains) antigen-specific T-cell receptors (TCRs), coded for by somatically recombined TCR genes. Each pathogen antigen is recognised by a small number of specific clones of T-cells, so that an individual animal contains a large, diverse repertoire of T-cell clones. T-cell responses to individual pathogens are idiosyncratic (Weng 2023); for example, in people infected with *Mycobacterium tuberculosis* individuals’ CD4^+^ T-cells recognised an average of 24 different epitopes, but individuals varied extensively in the epitopes that they recognised (Lindestam Arlehamn *et al*. 2016).

Next generation parallel sequencing of TCR-coding genes and transcripts has been used to investigate the TCR-coding sequences present among circulating T-cells in people and in laboratory mice. Theoretically, the TCR repertoire is enormous: in people potentially up to 10^15^ alpha and beta TCRs, though reduced to some 10^13^ after thymic selection, and then absolutely limited by the number of T-cells present in an individual, some 10^12^ in a person and 10^8^ in a laboratory mouse (Qi *et al*. 2014; Nikolich-Žugich *et al*. 2004). Sequence-based analyses are therefore only sampling a small fraction of the actual T-cells present, and so only sampling a small sub-fraction of the TCR repertoire (Laydon *et al*. 2015).

To date, most analyses of the TCR repertoire have been of people, with some study of laboratory mice. Many studies have confirmed that people and laboratory mice have large TCR repertoires. The clone size distribution of the repertoire is approximated well by a power law, with the vast majority of T-cell receptors found at very low frequency (typically observed once in a sample), but a proportion of large clones taking up a disproportionate part of the overall repertoire (Oakes *et al*. 2017; Sun *et al*. 2017). These distribution profiles can vary quantitatively between CD4^+^ and CD8^+^ cells, between naïve and memory cells, and as individuals age (Britanova *et al*. 2016; Oakes *et al*. 2017). Specifically, as people age there is a reduction in overall TCR diversity, driven in part by a decrease in the number of TCR-defined clones in naïve, but not in memory CD4^+^ cells, with similar changes in CD8^+^ cells, though they have a comparatively smaller TCR repertoire (Song *et al*. 2025; Weng 2023). The decrease in diversity of memory cells is driven by an increase in the relative abundance of some clones; indeed, in some individuals individual clones specific to a chronic virus such as Cytomegalovirus can occupy a significant proportion of the total repertoire (Lindau *et al*. 2019; Khan *et al*. 2002).

Here we sought to investigate the TCR repertoire of wild mice to complement other studies that have investigated wild mouse immune state. Animals’ exposure to pathogens and to other antigens drives the expansion of specific T cell clones, such that individuals may vary in their TCR repertoire because of differences in their lifetime of exposure, so that TCR repertoire may vary in mice of different ages. Male and female mice also live differently, with females living and breeding communally, but males being more solitary (Manning *et al*. 1995, Pocock *et al*. 2005), and this may affect exposure to infection such that there are sex differences in TCR repertoire. Therefore here we have investigated if, and how, the TCR repertoire varies among wild mice of different sex and age, and among those obtained from different sites. We present these data as a first analysis of wild mouse TCR repertories that is particularly relevant to eco-immunology, but also to laboratory mouse immunology.

## MATERIALS AND METHODS

### Animals

We live trapped mice, *Mus musculus domesticus* between May and September 2023 at two sites: (i) a pig farm in Nottinghamshire and (ii) a dairy farm on the Wirral peninsula, both in the UK. Mice were killed by cervical dislocation, sexed, and weighed. We aged mice by determining the mass of their dried eye lenses after Rowe *et al*. 1985; this method has been validated against mice of known age. We also analysed the repertories of 6 female CD-1 laboratory mice.

### Splenocytes, FACS, and isolation of RNA

From each mouse we aseptically isolated the spleen, determined its mass and then passed it through a 70 μm sieve to create a single-cell suspension. We resuspended the splenocytes in blood cell lysis buffer (Invitrogen) held at room temperature for 5 minutes, supplemented with an excess of phosphate buffered saline (PBS), sedimented the cells by centrifugation, washed them in PBS by resuspension and sedimentation, after which the cells were resuspend in 5 mL of staining buffer (Biolegend). We determined the number of splenocytes using a haemocytometer. We removed a portion of the cells as the FACS negative control. To the remainder we blocked Fc receptors using TruStain FcX™ PLUS on ice for 10 minutes and then stained the cells by addition of the following conjugated antibodies: anti-CD3-PE, anti-CD4-FITC, and CD8-APC for 20 minutes in the dark, after which the cells were washed twice in staining buffer. Immediately before FACS the cells were additionally stained with DAPI.

Samples were analysed and separated using a FACS Aria III Cell Sorter (BD Bioscience, Oxford, UK). The gating strategy was incorporated from dot plots with the following hierarchy: scatter gates (SSC/FCS) to identify mammalian cell clusters, distinct from debris with low particle size and granularity that were expected to be excluded from the gate. A singlet gate was applied (FSC-H / FSC-A) to remove doublets / multiplets that deviated from cluster linearity. From the singlet population, particles with negative signal toward the DAPI stain were considered membrane-intact and hence gated as live cells. The CD3^+^ population was used to identify T-cells, and finally CD4^+^ (negative to CD8) or CD8^+^ (negative to CD4) to detect and sort corresponding T-cell subsets (**Supplementary Material 1**). On-site data visualization, gating, and sort command were performed using FACS Diva software, version 8.1.

The resulting CD4^+^ and CD8^+^ sorted cells were concentrated by centrifugation, and the supernatant removed and RLT buffer (Qiagen) with 1 % (v/v) 2 mercaptoethanol added to the pellet of cells, and mixed by pipetting to form a lysate, which was then processed through a QIAshredder homogeniser spin column following the manufacturer’s methods, with the resultant RNA stored at −80°C.

### TCR sequencing and analysis

From the isolated RNA, TCR alpha and beta genes were sequenced using an established quantitative sequencing pipeline, which uses unique molecular barcodes to correct for PCR bias, and sequencing errors (Uddin *et al*. 2019; Oakes *et al*. 2017). The fastq files were processed and annotated using an in-house open source computational pipeline Decombinator V4 (Peacock *et al*. 2021; Decombinator 2024). We excluded any TCR nucleotide sequences that did not contain a CDR3 sequence.

### Data Analysis

Each TCR nucleotide sequence defines a different T-cell clone, where the number of occurrences of an identical nucleotide sequence is a measure of the abundance of that T-cell clone. Different TCR nucleotide sequences can code for the same TCR amino acid sequence. In the analyses that follow we therefore refer to, and differentiate, nucleotide-defined TCRs and amino acid-defined TCRs. Amino acid-defined TCR sequences were the predicted translation of the concatenation of the nucleotide sequences of the variable gene segment (V), joining gene segment (J) and CDR3 region.

We quality controlled the amino acid-defined TCR sequence data by examining the number of sequences obtained for alpha and beta chains of CD4^+^ and CD8^+^ cells and then excluding data for a chain where there were fewer than 5,000 TCR molecules that passed our quality control for that chain. We also examined the frequency distribution of amino acid sequences and fitted a power law distribution to it to generate an alpha coefficient, and then examined the relationship of this with wild mouse age. To test whether the frequency distribution of amino acid sequences for individual mice matched the power law distribution we used the Kolmogorov-Smirnov test with 500 simulations. Mice were classified into three categories based on the resulting p values: Fits power law (p > 0.1), Borderline fit to power law (0.1 > p > 0.05), Does not fit power law (p < 0.05). Results were summarised for each receptor type. The slope of the distribution did not vary with age.

We define TCR repertoire size as the sum of the number of occurrences of all amino acid-defined TCR sequences, separately for alpha and beta genes from CD4^+^ and CD8^+^ cells. We also calculated the number of unique amino acid-defined TCR sequences for each of the alpha and beta chains of CD4^+^ and CD8^+^ cells.

We calculated the richness of the TCR repertoire as the number of unique amino acid-defined TCRs as a proportion of the repertoire size, which corrects for the depth of sequencing of the samples. For the wild mice we tested for differences in the richness of the TCR repertoire between the alpha and beta chains of CD4^+^ and CD8^+^ cells using a gamma-distributed generalised linear model (GLM) with TCR richness as the response variable and RECEPTOR TYPE (*i.e.* CD4^+^ alpha, CD4^+^ beta, CD8^+^ alpha, CD8^+^ beta), SAMPLE SITE (Nottingham *vs*. Wirral), AGE, SEX, and an AGE*SEX interaction as explanatory variables. We calculated estimated marginal means (EMMs), which corrects for multiple comparisons, from the GLMs to test *post hoc* for differences between receptor types.

We calculated two diversity indices of the wild mouse TCRs: Shannon’s diversity index and Simpson’s diversity index. These metrics have recently been identified as being appropriate to quantify richness and evenness in TCR repertories (Mika *et al*. 2025). For both we down-sampled each mouse’s repertoire to that of the smallest, relevant repertoire (*i.e.* CD4^+^ alpha, CD4^+^ beta, CD8^+^ alpha, CD8^+^ beta) to account for differences in repertoire size between mice. To do this, we first replicated all TCR amino acid-defined sequences by their number of occurrences, then randomly sampled this list without replacement to the size of the smallest, relevant repertoire, and then re-compiled the TCR amino acid-defined sequences with their total number of occurrences. This down-sampling process was done separately for the two diversity measures.

Shannon’s diversity index was calculated as

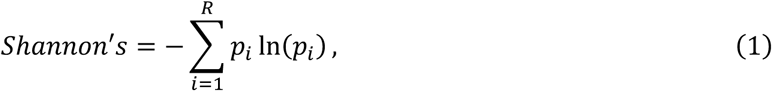

where *R* is the repertoire size, *p_i_* the proportional abundance of the *i^th^* TCR amino acid-defined sequence.

Simpson’s diversity index was calculated as

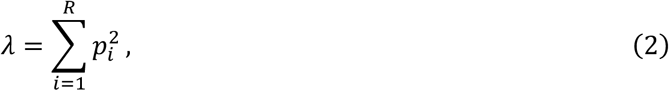

where *R* is the repertoire size and *p_i_* is the proportional abundance of the *i^th^* TCR amino acid-defined sequence. Analyses were conducted as for Shannon’s diversity index. We log transformed Simpson’s diversity index as

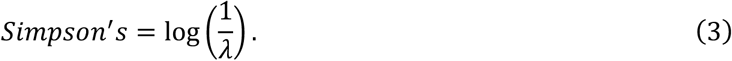

Thus, larger values indicate higher diversity.

We tested for differences in Shannon’s and Simpson’s diversity among the four receptor types (above) using a GLM (gaussian distribution) with Shannon or Simpson’s index as the response variable and RECEPTOR TYPE, SAMPLE SITE, AGE, SEX and an AGE*SEX interaction as explanatory variables. We calculated EMMs from the GLMs to test *post hoc* for differences between: (i) receptor types and (ii) diversity indices in male and female mice.

To investigate the degree of sharing of amino acid-defined TCR sequences between all pairwise combinations of the wild and laboratory mice, we first down-sampled each mouse’s TCR amino acid-defined repertoire to that of the smallest, relevant repertoire, as described above. We then compiled these repertoires into lists of unique amino acid-defined TCRs and performed pairwise comparisons between mice to determine the number of shared amino acid sequences between mouse pairs. From this we generated heatmaps of the degree of sharing, supplemented with dendrograms calculated from distance matrix data, which then arranged more similar individual mice together. We also examined these data using a principal coordinate analysis (PCoA) by calculating a re-scaled pairwise dissimilarity (with the most similar pair having a dissimilarity of 0), which was used in a weighted PCoA separately for alpha and beta chains of CD4^+^ and CD8^+^ cells.

To investigate whether mouse sex and age affected the extent to which mice shared amino acid-defined TCR sequences, we generated pairwise sharing data for all combinations of mice; we restricted this to only the Nottingham-derived mice to avoid any potentially effects of sample site. We then tested the effects of (i) pairwise mouse sex (same sex *vs.* different sex), (ii) pairwise difference in mouse age, and (iii) pairwise sum of mouse age on the number of pairwise shared amino acid-defined TCR sequences with a GLM (gaussian distribution). The response variable for the GLM was the number of pairwise shared amino acid-defined TCR sequences, while the explanatory variables were RECEPTOR TYPE, pairwise SEX DIFFERENCE, pairwise AGE DIFFERENCE, pairwise AGE SUM, a pairwise AGE DIFFERENCE*RECEPTOR TYPE interaction, and a pairwise AGE SUM*RECEPTOR TYPE interaction. We calculated slope estimates for the explanatory variables from the GLM using EMMs to test *post hoc* for differences between receptor types.

We conducted all data analysis in R (R Core Team 2025). Model diagnostics were carried out using the ‘DHARMa’ package (Hartig 2024). EMMs *post hoc* testing was carried out using the ‘emmeans’ package (Lenth 2025). PCoAs were conducted using the ‘vegan’ package (Oksanen *et al*. 2025). In addition, we used the ‘dplyr’ (Wickham *et al*. 2023), ‘stringr’ (Wickham 2023), ‘purrr’ (Wickham & Henry 2025), ‘tidyr’ (Wickham *et al*. 2024) ‘poweRlaw’ (Gillespie 2015), ‘here’ (Müller 2020), and ‘modeest’ (Poncet, 2019) packages. Plots were generated in R using packages ggplot2 (Wickham 2016), ‘gplots’ (Warnes *et al*. 2024), ‘showtext’ (Qiu 2024), ‘patchwork’ (Pederson 2024), ‘ggpubr’ (Kassambara 2023) and ‘ggbeeswarm’ (Clarke *et al*. 2023).

## RESULTS

### Animals

We caught 69 mice (30 male, 39 female), 65 from the Nottingham site and four from the Wirral site. The average age of the wild mice was 13 weeks (SD 10, median 9, range 2 – 49); average body mass was 15.5 g (SD 4, median 11, range 7 – 27). Among the wild mice, the spleens had an average mass of 53 mg (SD 33, range 10 – 170), yielding an average of 16 million splenocytes (SD 13, range 8.7 x 10^4^ – 5.1 x 10^7^). There was a positive relationship between mouse age and spleen mass (R^2^ = 0.3, p < 0.0001) and number of splenocytes (R^2^ = 0.13, p < 0.0025), and between spleen mass and number of splenocytes (R^2^ = 0.49; p, 0.0001) (**Supplementary Material 2**). Previous comparisons of wild and laboratory mice have found that wild mice are smaller than laboratory mice, and that their spleens are absolutely smaller (approximately one third of the mass) and with commensurate fewer cells, but also proportionally smaller (compared with body mass), compared with those of laboratory mice (Abolins *et al*. 2017).

The FACS of splenocytes resulted in an average of 8 million CD3^+^ T-cells per wild mouse (SD 8 million, range 1.1 x 10^4^ – 4.9 x 10^7^). There were proportionally more CD4^+^ cells than CD8^+^ cells, with an average of 62 % (SD 12, range 34 – 82), giving an average CD4^+^ : CD8^+^ ratio of 2.4 (SD 1.4, range 0.2 - 6.9). For four female mice from the Nottingham site insufficient cells were collected for TCR sequence analysis, refining our sample size to 65 (30 male, 35 female) wild mice and 6 lab mice.

### TCR sequences

To quality control the data we examined the number of alpha and beta chain coding sequences for CD4^+^ and CD8^+^ cells separately, and excluded the data for one cell type if there were fewer than 5,000 TCR molecules of either the alpha or beta chain, which resulted in the exclusion of 9 and 13 mice, respectively. This resulted in final sample sizes of 62 (56 wild, 6 lab) for CD4^+^ alpha and beta, and 58 (52 wild, 6 lab) CD8^+^ alpha and beta chain TCR sequences that were analysed further.

After our data quality control the minimum TCR repertoire size among the wild mice was 54,908, 58,772, 66,881 and 64,224, and for the laboratory mice the minimum 68,637, 79,052, 49,379 and 75,640 for CD4^+^ alpha and beta and CD8^+^ alpha and beta chains, respectively. We examined the frequency distribution of the amino acid-defined TCR sequences, which showed that many of these followed a power law distribution, which has frequently been observed on other TCR datasets (**Supplementary Material 3**).

### TCR repertoire richness and diversity

We investigated the richness of the TCR repertoire, finding that among the wild mice beta chain richness was greater than that of the alpha chain, for both within and between CD4^+^ and CD8^+^ cell comparisons (**Supplementary Material 4**). The comparatively larger repertoire of beta chains, observed previously in human T-cells (Oakes *et al*. 2017), occurs because they are encoded by recombination of V, D and J genes, unlike the V and J recombination of alpha chain coding genes.

There was no significant effect of wild mouse AGE, SAMPLE SITE, SEX, or AGE*SEX on the TCR repertoire richness (all |z| < 1.2, p > 0.26; **Supplementary Material 4**). However, the TCR diversity measured by Shannon’s index (after down-sampling each mouse’s repertoire to that of the smallest, relevant repertoire) differed significantly between the CD4^+^ beta and the CD8^+^ alpha chains (**Figure 1; Supplementary Material 5**), and there were significant effects of SEX (t = 2.79, p = 0.0058), SAMPLE SITE (t = 2.76, p = 0.0063), SEX*AGE (t = −2.07, p = 0.0396), and a marginally non-significant effect of AGE (t = −1.78, p = 0.0762) with older mice tending to have a lower diversity. The difference between Shannon’s index of female and male mice was marginally non-significant (effect size = - 0.0996, p = 0.0859). We obtained broadly similar results using Simpson’s diversity index though with more significant differences in diversity between receptor types (**Supplementary Material 6**), significant effects of SEX (t = 2.02, p = 0.0452), and AGE (t = −3.97, p = 9.91 x 10^-5^), with older mice having lower diversity (**Supplementary Material 7**). The difference between Simpson’s index of female and male mice was also marginally non-significant (effect size = −0.187, p = 0.0884).

**Figure 1.**
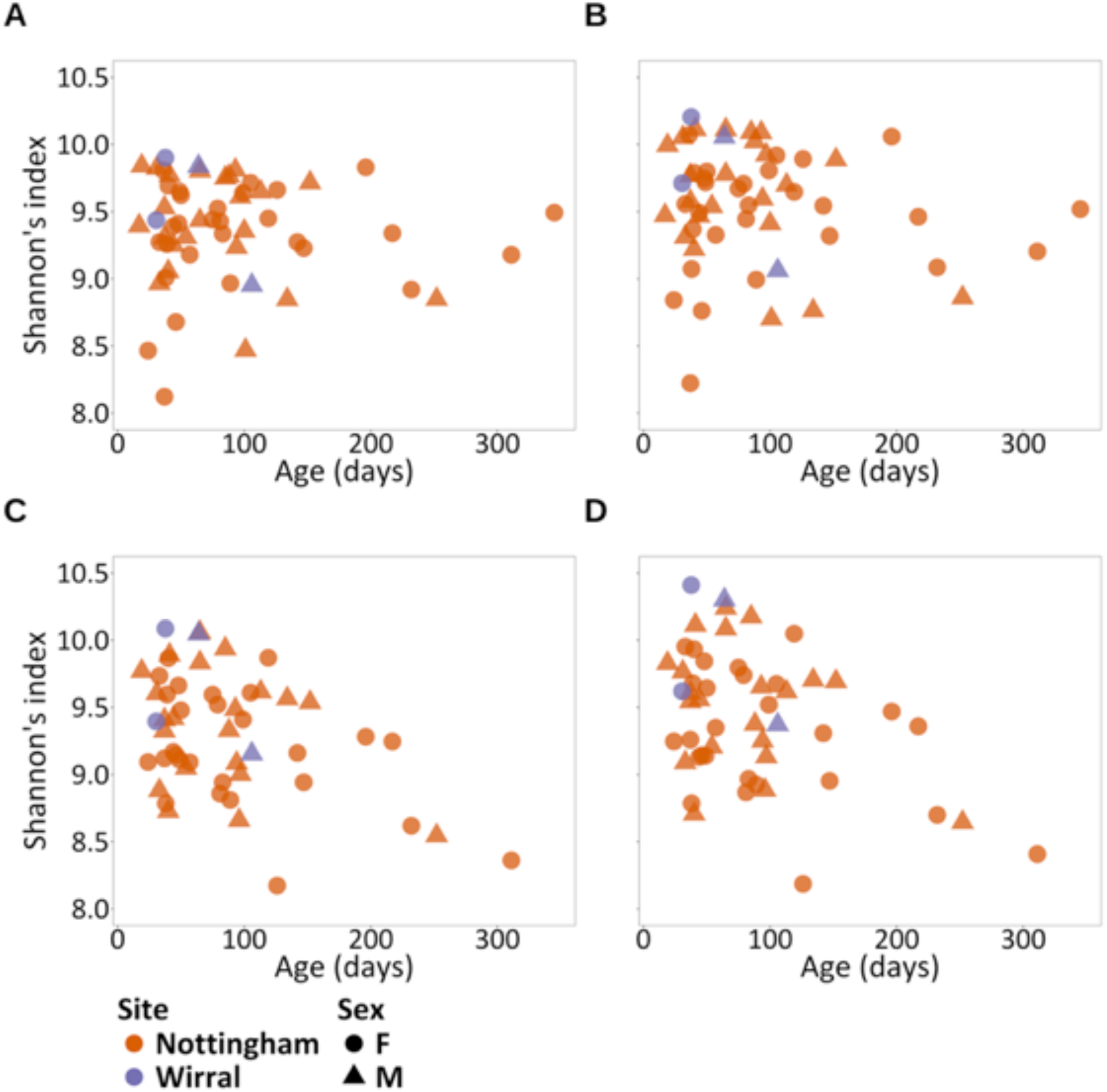
Alpha diversity measured by Shannon’s index of wild mouse, amino acid-defined TCRs for (A) CD4^+^ alpha, (B) CD4^+^ beta, (C) CD8^+^ alpha and (D) CD8^+^ beta cells.

### TCR networks

We investigated how much different wild and laboratory mice shared the same amino acid-defined TCR sequence, which we did by counting the number of common sequences between every pairwise combination of mice. This showed that there was considerable sharing of TCR sequences, with this greatest for alpha chains (5.9 and 5.2 % for CD4^+^ and CD8^+^, respectively) than for beta chains (2.9 and 2.2 % for CD4^+^ and CD8^+^, respectively) (**Figure 2**; **Supplementary Material 8**). We asked how many TCR sequences were public, which we defined as those shared among ≥75 % of the mice (both wild and laboratory). We found that this ranged from 157 (for CD8^+^ beta) to 1,113 (for CD4^+^ alpha), but that this was always less than 0.25 % of unique TCRs (**Supplementary Material 8**). Previous studies of public TCR sequences in 28 laboratory mice found that ∼0.5 % of sequences were shared among 75 % of them (Madi *et al*. 2014). A PCoA analysis showed that there were differences in the sharing of sequences among mice from the different wild sampling sites, and between wild mice and laboratory mice (**Supplementary Material 9**).

**Figure 2.**
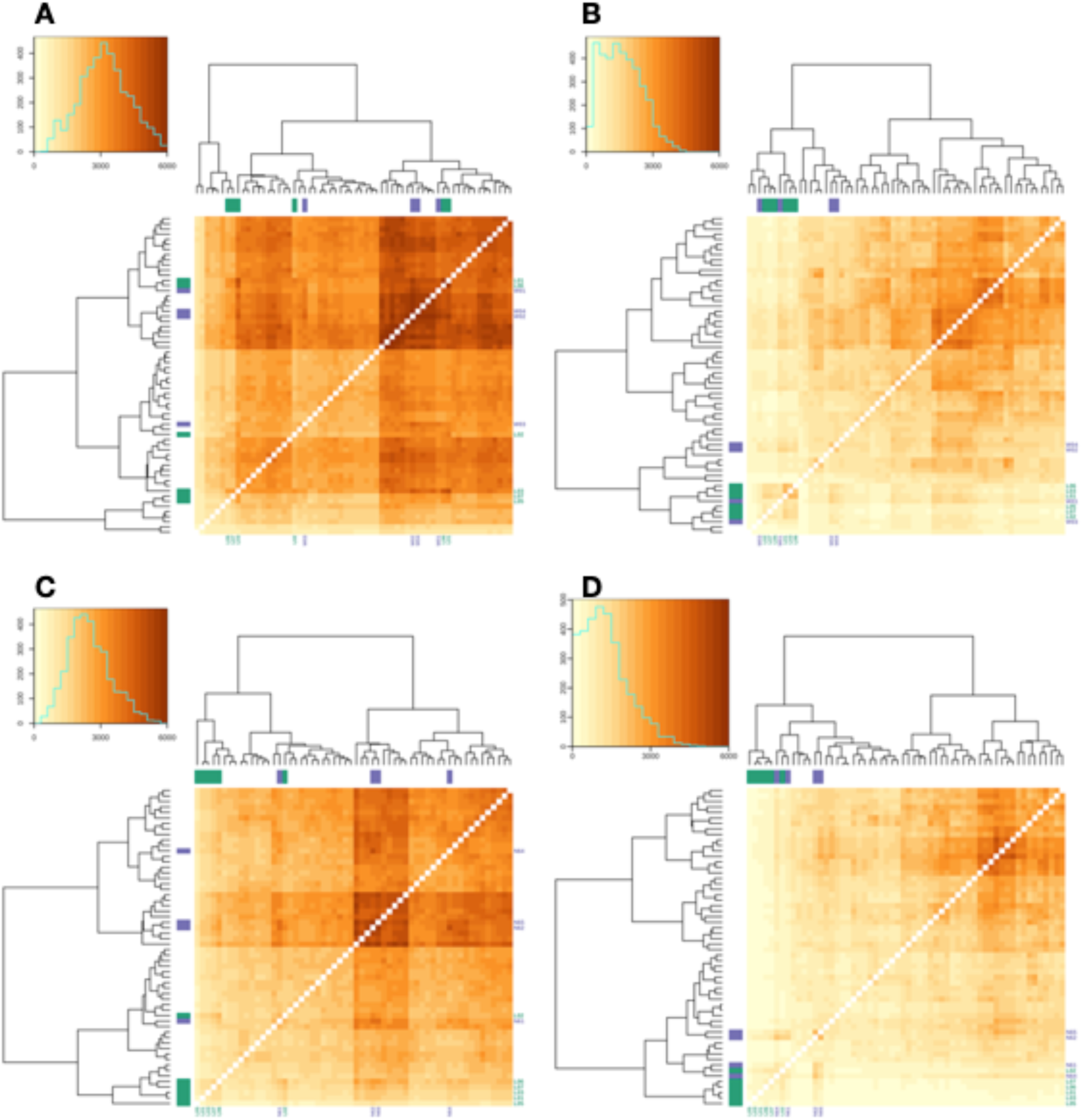
The sharing of amino acid-defined TCR sequences among wild and laboratory mice, with the colour scale showing the number of shared sequences for (A) CD4^+^ alpha, (B) CD4^+^ beta, (C) CD8^+^ alpha, (D) CD8^+^ beta. Against the plots, purple indicates wild mice from the Wirral site and green laboratory mice; mice where no colour is shown are wild mice from the Nottingham site. In all plots the brown scale is the same. In each, the insert plot shows a frequency distribution of the number of shared sequences; note that the y-axis is a different scale in these four plots, though the x-axis is consistent.

We investigated whether mouse sex and age affected the extent to which mice shared amino acid-defined TCR sequences. Mindful of the differences in sharing among mice from the different wild sites, we focussed this analysis on only the mice from the Nottingham site, and used a GLM that analysed the number of shared sequences by (i) same sex *vs.* different sex pairs, (ii) sum of age of mouse pairs, and (ii) difference in age of mouse pairs. We used these sum and difference in age metrics to see if individuals’ TCR repertoires converged or diverged as mice age. This showed that overall there was no difference in the number of shared TCR sequences between same sex and different sex pairs (t = 0.398, p = 0.69), and that the sum of mouse pair age was not significant (t = 0.180, p = 0.47), nor for the difference in mouse pair age (t = −0.541, p = 0.122), (**Figure 3; Supplementary Material 10**). However, *post* hoc analysis of the four different receptor types showed that for CD8^+^ alpha and beta receptors there were consistent negative relationships of the number of shared sequences and the sum or difference of age (**Supplementary Material 11**). This was also observed for CD4+ beta chains, but not alpha.

**Figure 3.**
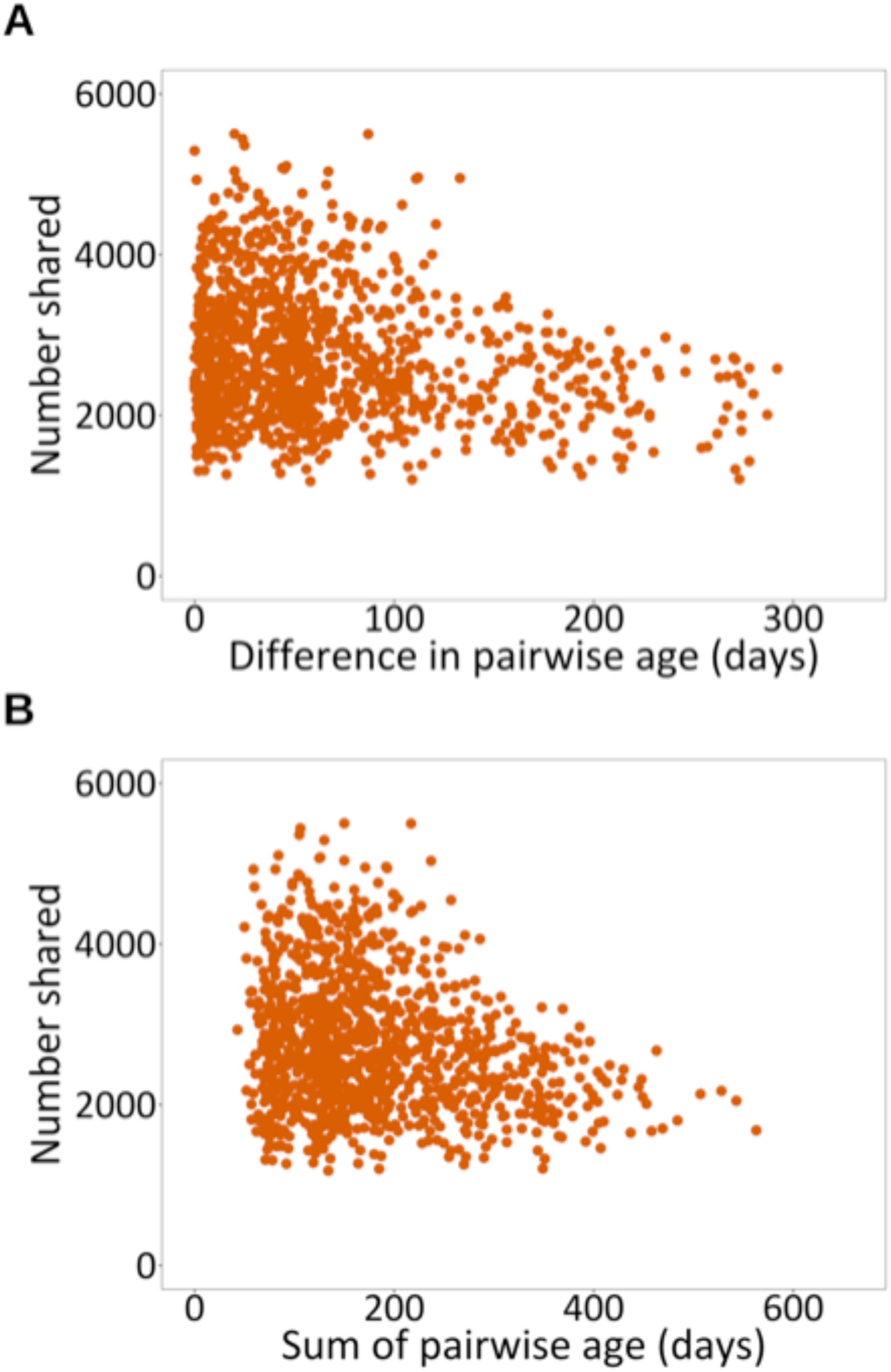
The number of shared amino acid-defined TCR sequences for CD8^+^ alpha between all pairwise combinations of wild mice from the Nottingham site against (A) the difference in pairwise age and (B) the sum of the pairwise age, where age is in days. The results for CD4^+^ alpha and beta, and CD8^+^ beta are shown in **Supplementary Material 10**.

## DISCUSSION

The aim of this work was to describe – for the first time, as far as we are aware – the TCR repertoire of wild mice and to investigate if, and how, individuals’ age, sex, and the site from which they were obtained affected their TCR repertoire. This work is directly relevant to the fields of eco-immunology and mouse immunology more generally. We successfully sampled wild mice and obtained TCR sequences from CD4^+^ and CD8^+^ cells. Despite wild mice being smaller than laboratory mice and having absolutely and proportionally smaller spleens with fewer splenocytes (Abolins *et al*. 2017) we successfully obtained sufficient CD4^+^ and CD8^+^ cells from 65 wild mice for TCR sequence analysis.

Wild mice have large TCR repertoires and their richness was not affected by individuals’ sex, age, or the site from which the mice were obtained. Richness (which is the normalised number of unique TCRs in a repertoire) measures the breadth of the TCR repertoire. We hypothesise, therefore, that both male and female mice maintain the capacity to respond to a wide range of antigens in both CD4^+^ and CD8^+^ T-cell compartments throughout their natural lifespan in the wild. This contrasts with some studies in humans that report declining richness with age (Weng, 2023). These differences between wild mice and people may reflect some immunological differences between wild mouse and human populations. But in addition to this (i) our sampling of wild mice will have under-sampled very young mice (which typically will not leave their nests), so that we may not have been able to account for TCR richness in very young individuals, and (ii) the maximum lifespan of wild mice is not equivalent to that of elderly people included in these studies and / or the relative rarity of old mice limits our statistical power.

In contrast to TCR repertoire richness, measures of diversity (such as Shannon and Simpson’s indices) assess both the number of different TCRs and their relative abundance of those, such that when there are more abundant clones then the diversity is lower. The two diversity indices give slightly different results, but concur in showing an effect of sex on TCR diversity, and generally agree in showing an effect of age; Shannon’s index also shows an effect of an interaction between age and sex. In the context of constant richness, decreased diversity must reflect an increase in the abundance of some clones, which we hypothesise may arise from chronic exposure to a limited set of infections prevalent in wild mouse populations. An influence of sex on diversity has been reported previously in humans, with greater diversity in females (Trofimov *et al*. 2022). The mechanism behind these effects remains unclear, but could reflect hormonal differences leading to differences in thymic selection (Trofimov *et al*. 2022). Decreased diversity with age has also been described repeatedly in humans (Britanova *et al*. 2016). In wild mice the effects of sex and age may reflect differences among mice in their exposure to infection, with higher diversity a consequence of less exposure to infection. Thus, young mice will not yet have had time to experience many infections unlike older mice. Male house mice may have more limited social interactions because they typically disperse from the site of their birth (Pocock *et al*. 2005) leading to comparatively less intense or less chronic exposure to infection in their environment. In contrast, females breed communally (Manning *et al*. 1995), which may expose them to greater levels of infection and antigenic challenge. Antigenic sources beyond infection – including self antigens and tumours – will also result in T-cell responses, and these may contribute to the age and sex effects on TCR diversity. However, the antigenic assault from infection is particularly strong in wild animals, especially compared to laboratory animals, and we suspect likely to be the major antigen source across wild animal populations. Taken together, our results suggest that wild mice maintain a broad TCR repertoire that allows them to react to a breadth of antigens but, particularly in older, female mice, this is evidence of strong, life-long immune responses to a limited set of persistent infections in their environment.

We found that individual mice shared a considerable number of TCR sequences; specifically *c.*5 % of alpha chains and *c.*2 % of beta chains. The differences in the amount of TCR sharing between alpha and beta chains is, presumably, because beta chains are comparatively more diverse, making the chance of them being shared less likely compared with alpha chains. We found a pattern of greater sharing between mice from within the same group (from Nottingham, from the Wirral, or laboratory mice) than between groups, which is suggestive of diverse antigenic experience of mice – leading to different TCR sequences – in the different environments of the groups of mice. This is consistent with previous work that found immunological differences between wild mice from different sample sites (Abolins *et al*. 2018).

We also found that wild mouse age affects the number CD8^+^ TCR sequences that are shared among mice. We suggest that this occurs because as mice age they become more idiosyncratic in their amino acid-defined TCR sequences, so that older mice share fewer sequences with younger mice (the effect of the difference in mouse age) and that older mice share fewer sequences with other, older mice (the effect of the sum of mouse age). Idiosyncratic responses of laboratory mice to protein antigens and viral infection has been reported previously (Sun *et al*. 2017; Milighetti *et al*. 2023). Study of wild mouse (*Mus musculus* and *Apodemus sylvaticus*) gut microbiomes show that they differ significantly among individuals (Marsh *et al*. 2022; Hanski *et al*. 2025), and this may result in idiosyncratic antigenic experience in the gut, which may contribute to the individuality of wild mouse TCRs that we have observed.

Our study has a number of limitations. One, we obtained our samples from live-trapped mice, and this method does not effectively sample very young animals, so that these are under-represented in our study. Two, we caught wild mice at only two sample sites, and most of the caught mice came from one site. This did not give us much statistical power to investigate how TCR repertoires may vary among individuals from different sample sites. This could be addressed by further work in the future, which could be extended to analyse both spatial and temporal aspects of TCR repertoire diversity. Three, we analysed only a small number of control, laboratory mice. Surprisingly, given the central role of laboratory mouse models in the development of modern immunology, there appear to be a dearth of publicly available data on the TCR repertoires of the common strains of laboratory mice. We have therefore been unable to make a comprehensive comparison of TCR repertoires between wild and laboratory mice. Given the already established differences in immune state between wild and laboratory mice, a future, in-depth comparison between the TCR repertoires of wild and laboratory mice would be of interest.

Immunological analyses of non-standard model species is often hampered by the paucity of immunological reagents that are available. However, because the TCR repertoire can be investigated using sequence-based analyses this approach can be widely applied to a whole range of species. In so doing, this may allow a broad, comparative study of wild animals’ TCR repertoires.

In summary, here we describe the TCR repertoire of wild mice and investigated how this is affected by individuals age, sex, and site of sampling. Our analyses of the wild mouse TCR repertoire finds some patterns that are similar to human TCR repertories, though we find differences between wild male and female mice that may reflect their different lives and exposure to infection.

## CONFLICT OF INTEREST DISCLOSURE

The authors have no conflicts of interest.

## CRediT AUTHORSHIP CONTRIBUTION STATEMENT

**Jacob Cohen**: Formal analysis, Validation, Visualization, Writing – original draft. **Simon Hunter-Barnett**: Data curation, Investigation, Project administration, Validation, Writing – review & editing. **Gayathri Nageswaran**: Data curation, Methodology. **Suzanne Byrne:** Data curation, Methodology. **Gemma Freeman**: Formal analysis. **Matthew Cowley**: Formal analysis, Writing – review & editing. **Benny Chain**: Conceptualization, Formal analysis, Funding acquisition, Project administration, Supervision, Writing – review & editing. **Mark Viney**: Conceptualization, Formal analysis, Funding acquisition, Project administration, Supervision, Writing – original draft. the paper.

## SUPPLEMENTARY MATERIAL

The Supplementary Material contains 11 items in a single file.

## DATA AVAILABILITY STATEMENT

The datasets generated for this study can be found in the Sequence Read Archive [www.ncbi.nlm.nih.gov/sra] as SUB15234028 / PRJNA1249624 and our processed repertories and mouse metadata are in Zenodo at https://doi.org/10.5281/zenodo.17241409. Code used to analyse data for this study can be found at https://github.com/jacohen1/Wild-mouse-TCR.

## FUNDING

This work was funded by a NERC Exploring the Frontiers grant, NE/X010295/1.

## ACKNOWLEDGMENTS

We would like to thank landowners for access; in Liverpool the staff of the Biomedical Services Unit, and Chris Law of the Cell Sorting Facility for their help.

---

**Supplementary Material 1.**
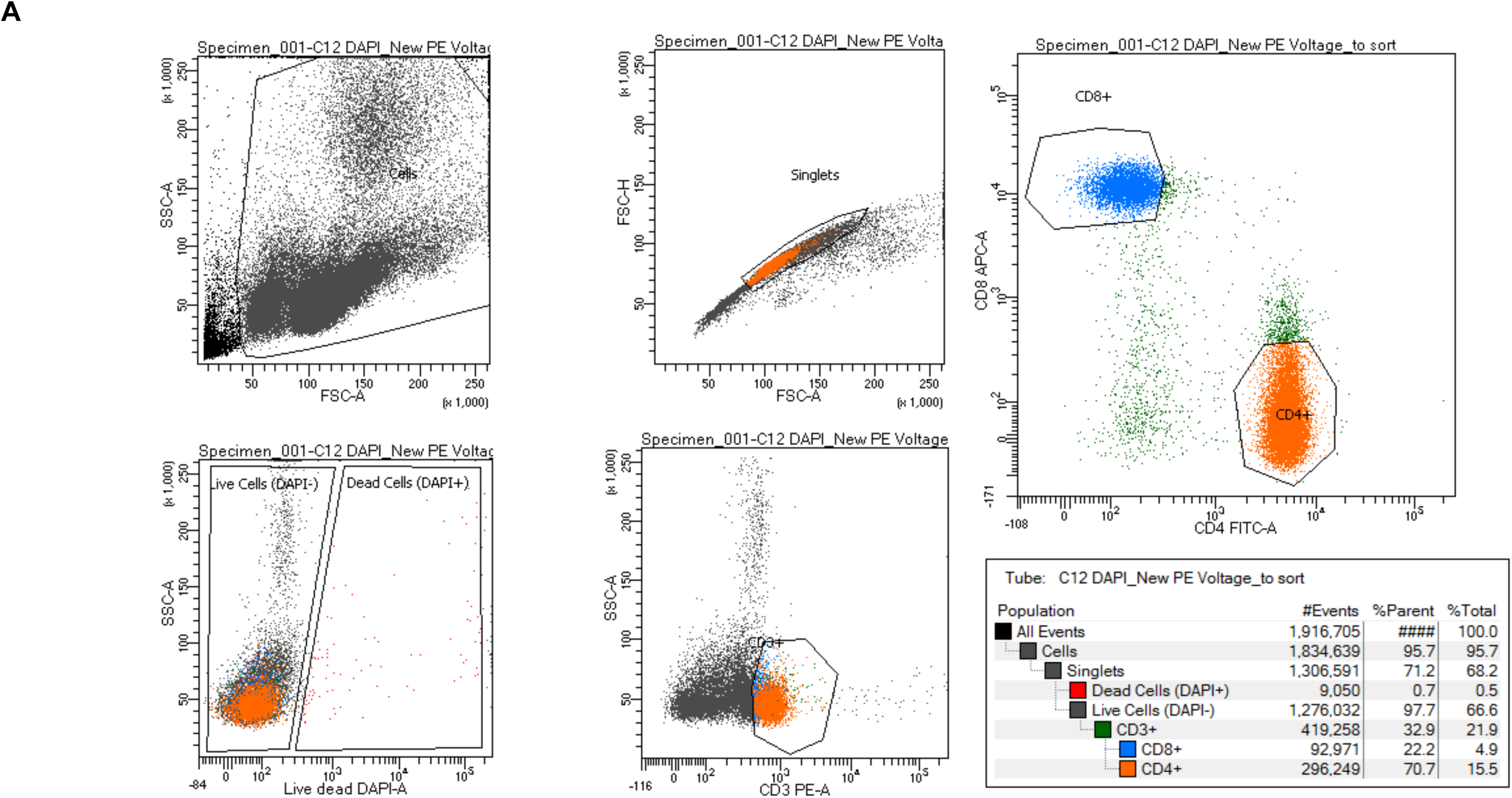
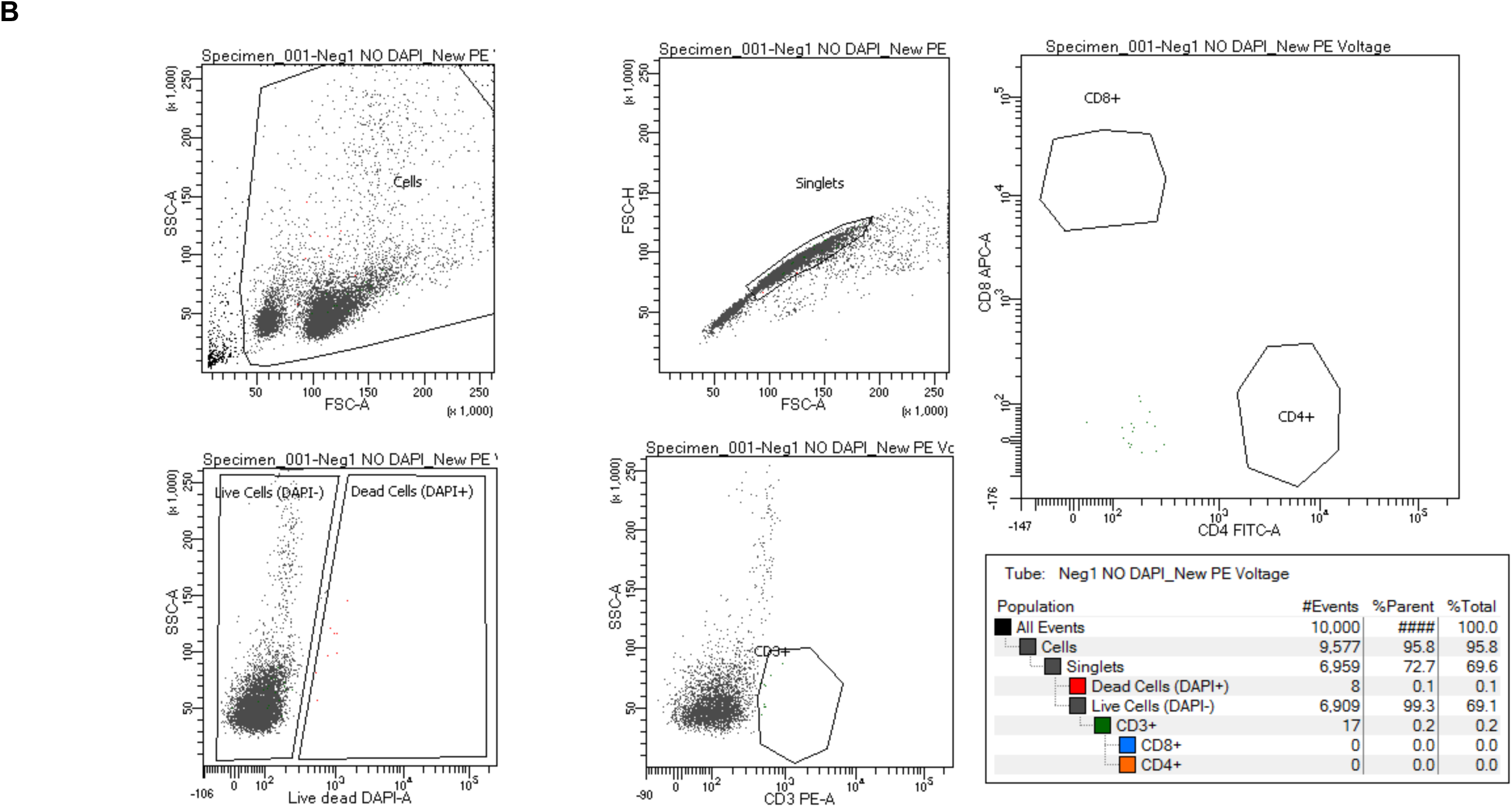
Example FACS plots of a mouse splenocyte sample where (A) cells were stained with DAPI, CD3-PE, CD4-FITC, and CD8-APC and (B) where for the same sample DAPI, CD3-PE, CD4-FITC, and CD8-APC were omitted from the staining protocol. For each, the gating hierarchy are plotted sequentially as (i) FSC-A *vs*. SSC-A, (ii) FSC-A *vs.* FSC-H, (iii) DAPI *vs.* SSC, (iv) CD3-PE *vs.* SSC, (v) CD4-FITC *vs.* CD8-APC, and the subsequent CD3^+^ CD4^+^ and CD3^+^ CD8^+^ were defined from dot plots of pair-matched negative controls (no antibody staining) from each individual (B), where maximum tolerance to false positive was <0.3 %.

**Supplementary Material 2.**
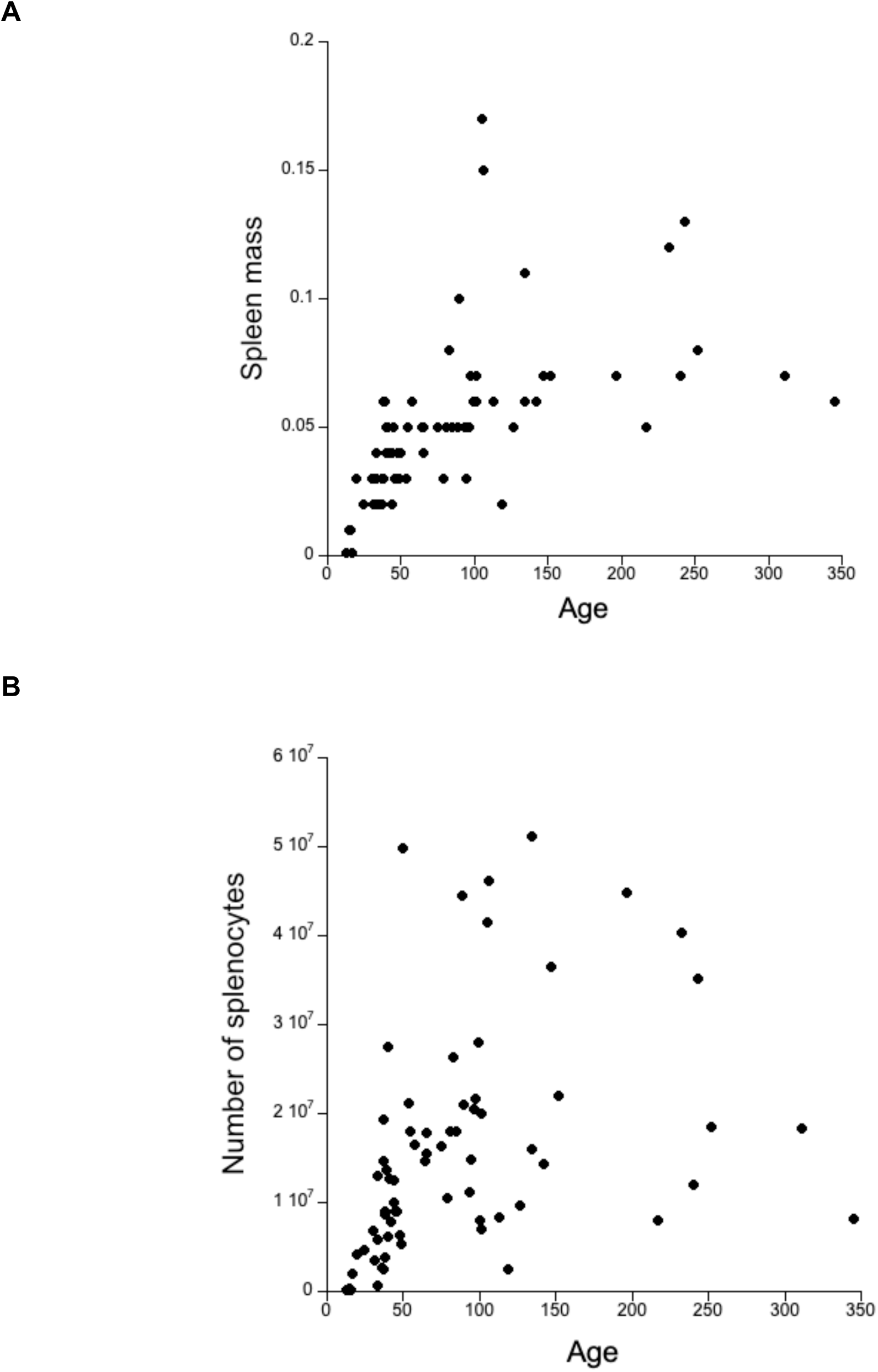
Mouse age in days and (A) spleen mass in grammes (Spleen mass = 0.03 + 0.0002.Age, R^2^ = 0.3, p < 0.0001) and (B) number of splenocytes (Number of splenocytes = 1.02 x 10^6^ + 6.37 x 10^4^.Age,R^2^ = 0.13, p < 0.0025).

**Supplementary Material 3.**
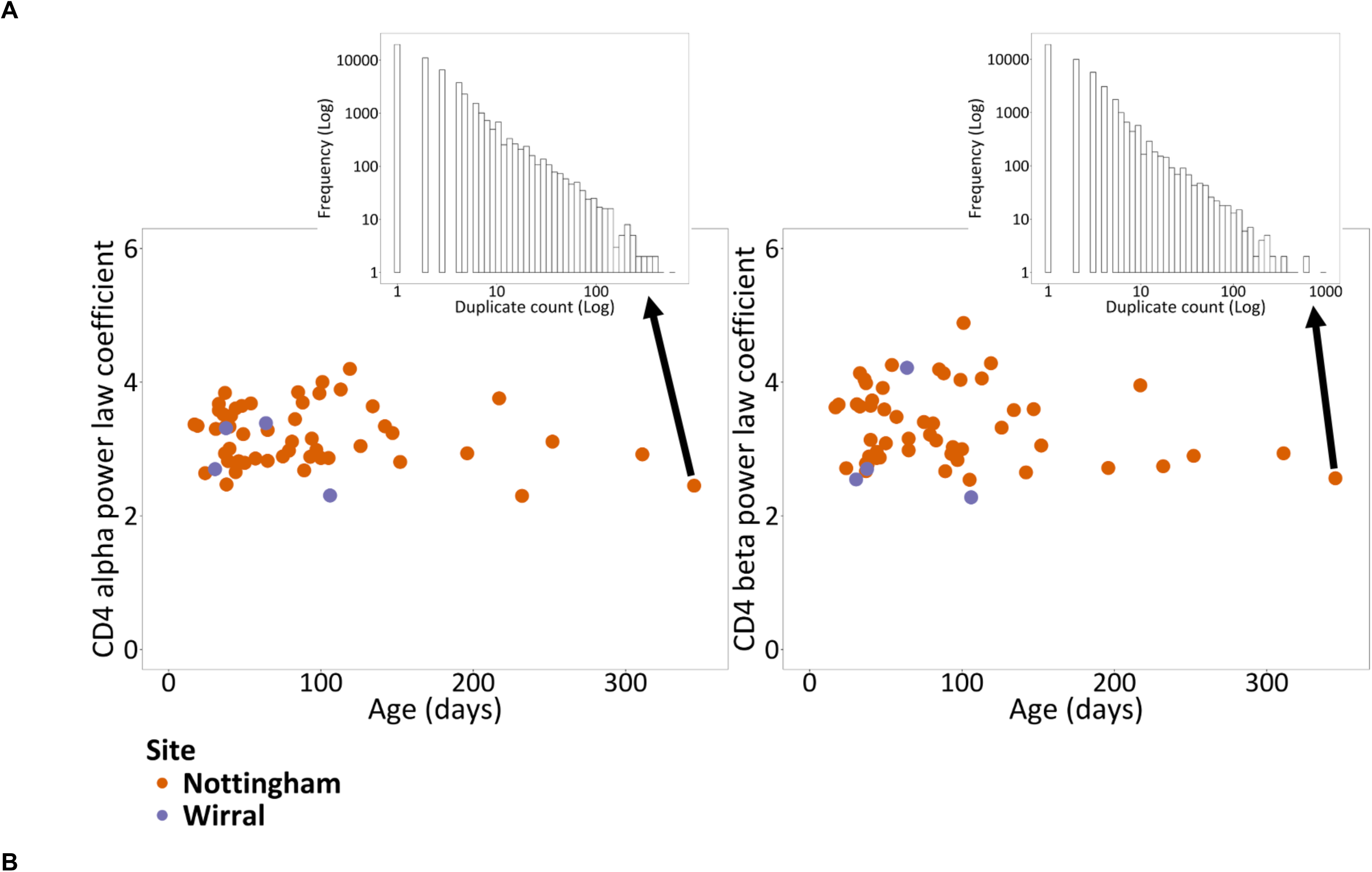
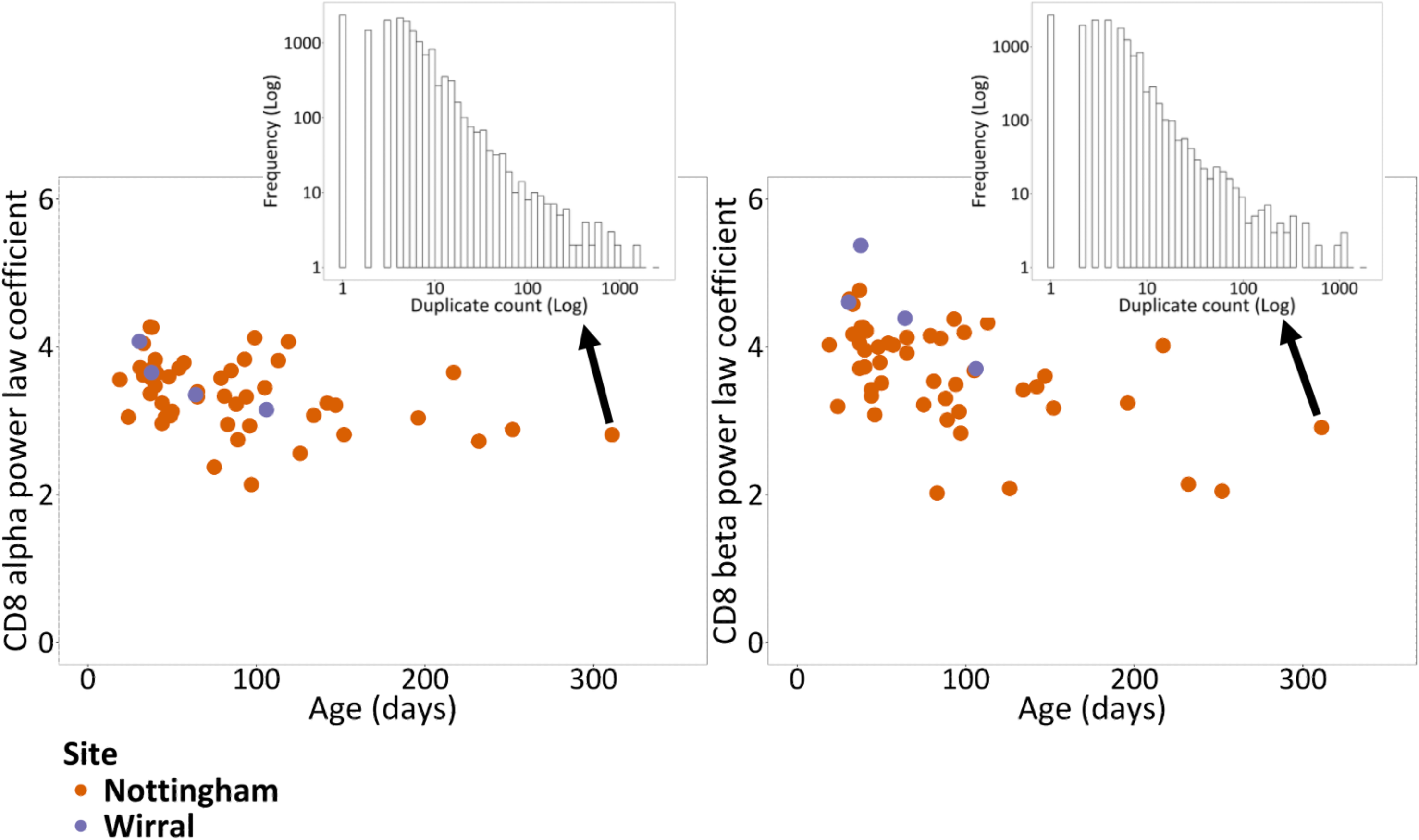
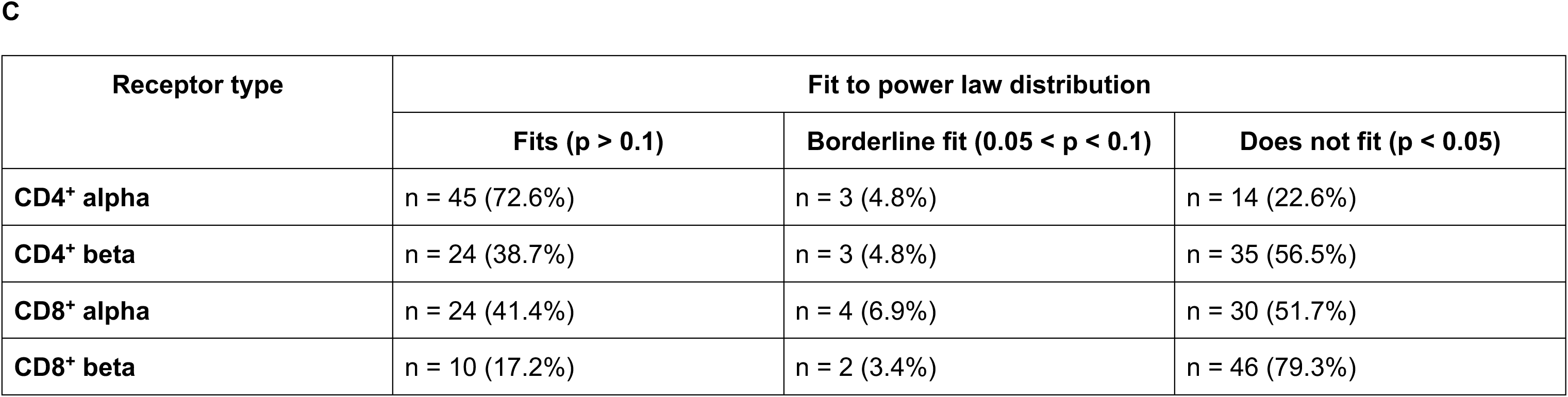
The power law coefficients of the frequency distribution of amino acid-defined TCR sequences for wild mice from the two sample sites for (A) CD4^+^ and (B) CD8^+^ cells; within each an example Log Frequency *vs.* Log Number of amino acid-defined sequences is shown. (C) Result of Kolmogorov-Smirnov test with 500 bootstraps to test that the data follows a power law distribution, showing the number of mice in each category.

**Supplementary Material 4.**
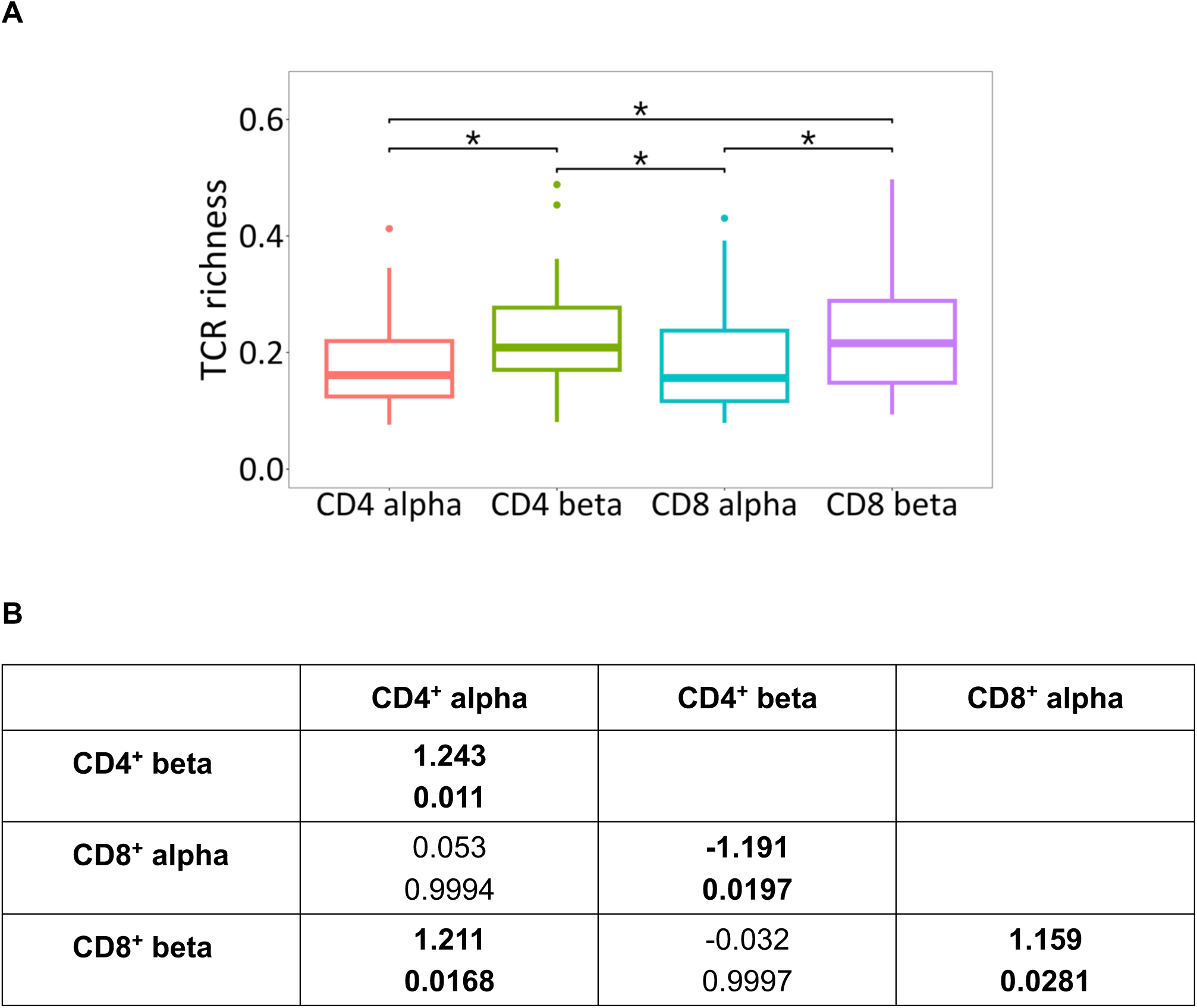
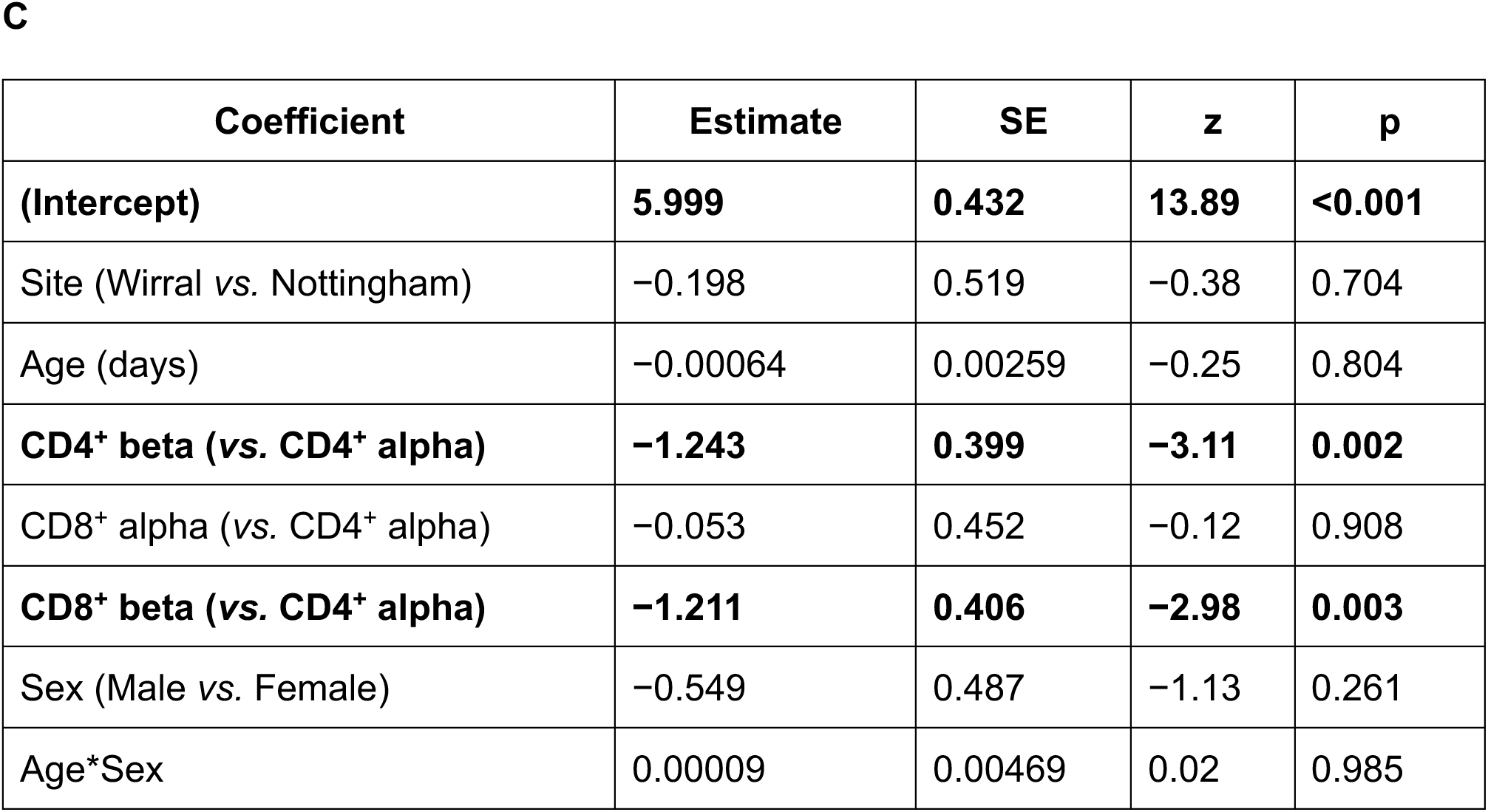
(A) Wild mouse TCR richness (calculated as the number of unique amino acid-defined TCRs as a proportion of the total number of amino acid-defined TCR sequences); the whiskers are 1.5x the inter-quartile range; * is p < 0.05. (B) The effect sizes (top value) and p values (bottom value) for pairwise comparisons of the marginal means estimated from a GLM of the form VALUE ∼ RECEPTOR TYPE + SAMPLE SITE + AGE + SEX; p < 0.05 are in bold. (C) Summary of wild mouse TCR richness GLM. Estimates are shown relative to reference levels (CD4^+^ alpha, Nottingham, Female). Coefficients with p < 0.05 are in bold. GLM details were: *n* = 216; residual df = 208; null deviance = 37.40; residual deviance = 33.61; AIC = −495.5.

**Supplementary Material 5.**
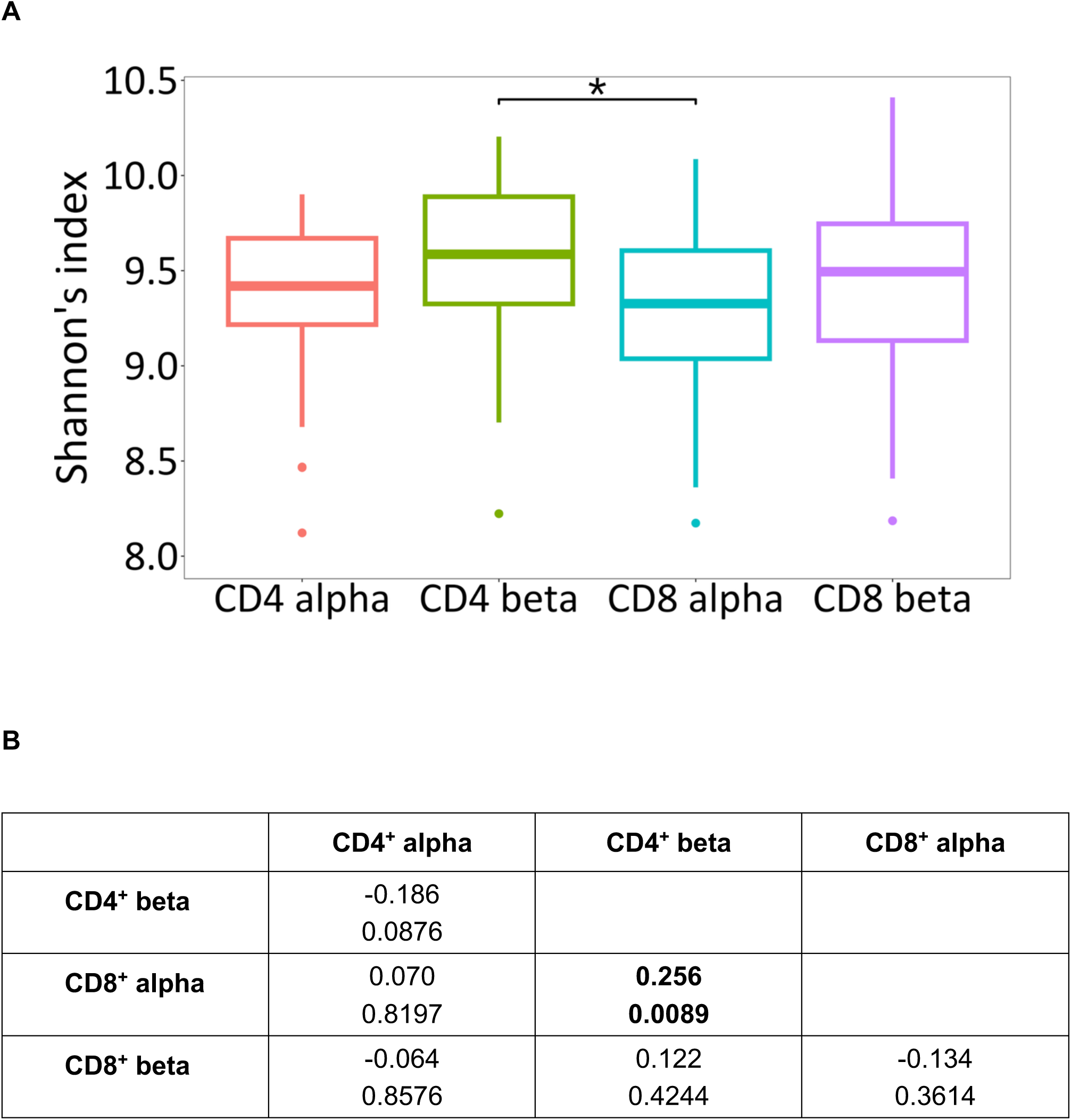
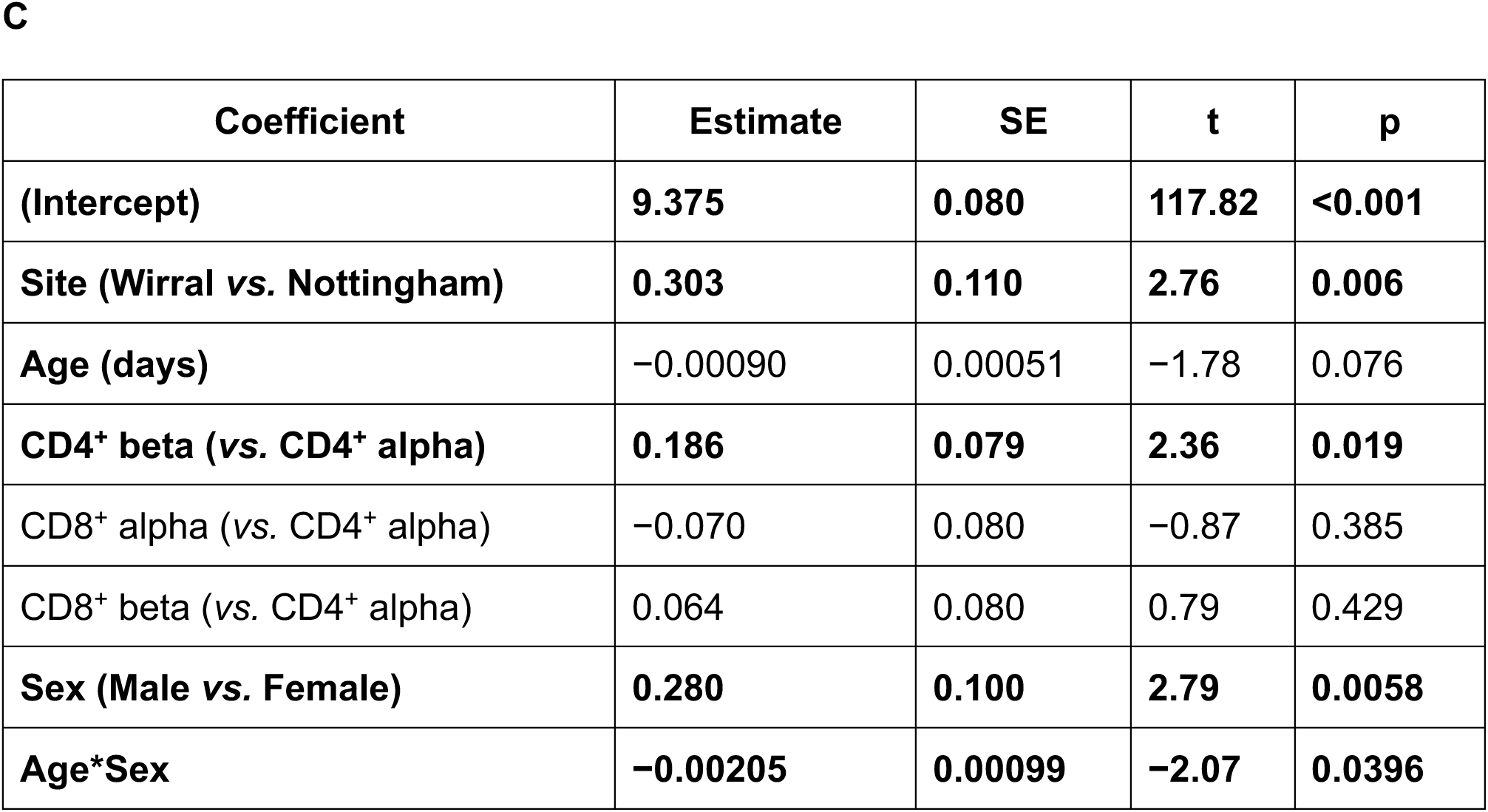
(A) Wild mouse amino acid-defined TCR alpha diversity using Shannon’s index; the whiskers are 1.5x the inter-quartile range; * is p < 0.05. (B) The effect sizes (top value) and p values (bottom value) for pairwise comparisons of the marginal means estimated from a GLM of the form VALUE ∼ RECEPTOR TYPE + SAMPLE SITE + AGE + SEX + AGE*SEX; p < 0.05 are in bold. (C) Summary of Shannon’s diversity index GLM. Estimates are shown relative to reference levels (CD4^+^ alpha, Nottingham, Female). Coefficients with p < 0.05 are in bold. GLM details were: *n* = 216; residual df = 208; null deviance = 43.23; residual deviance = 36.10; AIC = 244.5.

**Supplementary Material 6.**
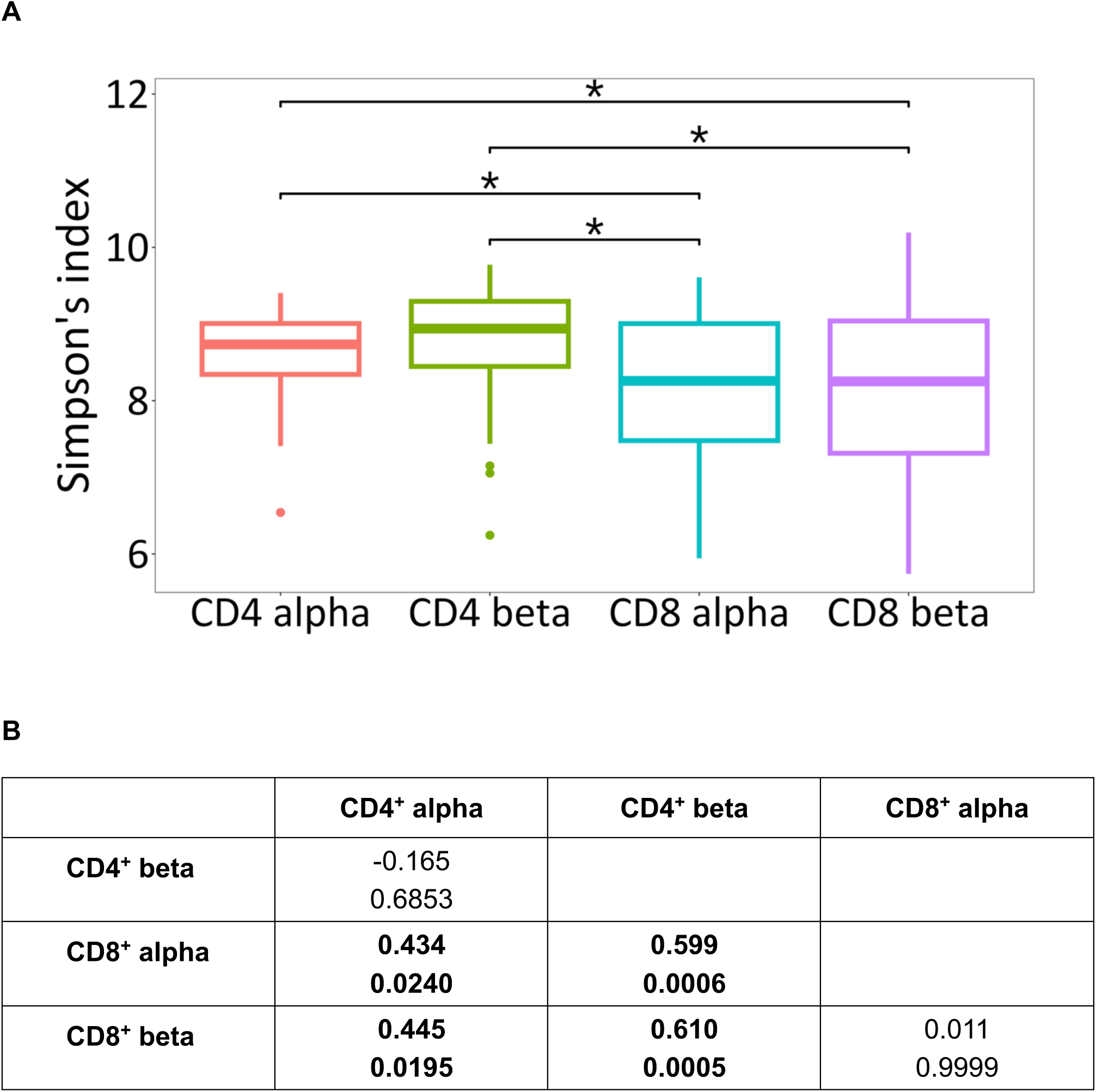
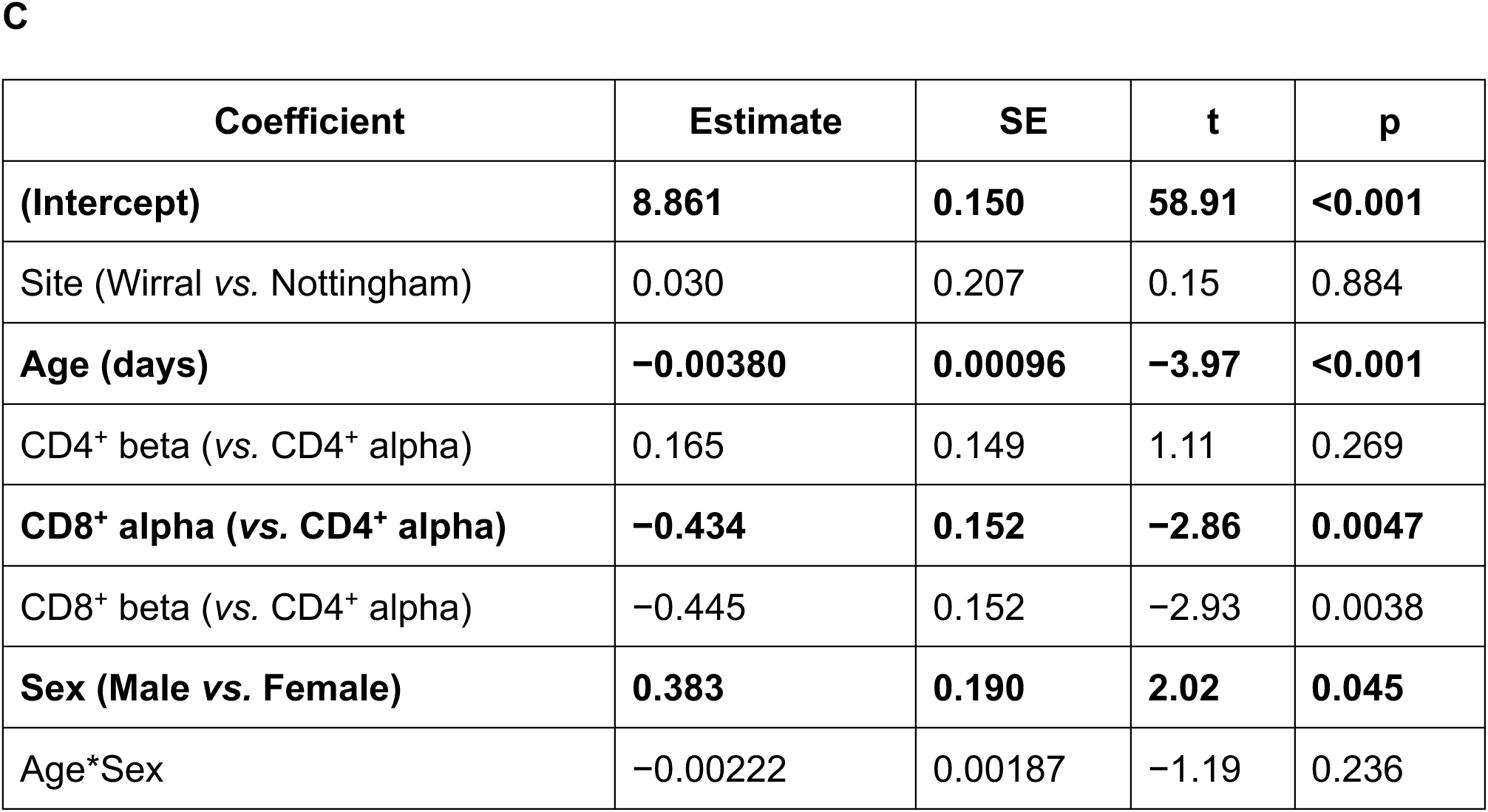
(A) Wild mouse amino acid-defined TCR alpha diversity using Simpson’s index; the whiskers are 1.5x the inter-quartile range; * is p < 0.05. (B) The effect sizes (top value) and p values (bottom value) for pairwise comparisons of the marginal means estimated from a GLM of the form VALUE ∼ RECEPTOR TYPE + SAMPLE SITE + AGE + SEX + AGE*SEX; p < 0.05 are in bold. (C) Summary of Simpson’s diversity index GLM. Estimates are shown relative to reference levels (CD4^+^ alpha, Nottingham, Female). Coefficients with p < 0.05 are in bold. GLM details were: *n* = 216; residual df = 208; null deviance = 166.8; residual deviance = 129.0; AIC = 519.6.

**Supplementary Material 7.**
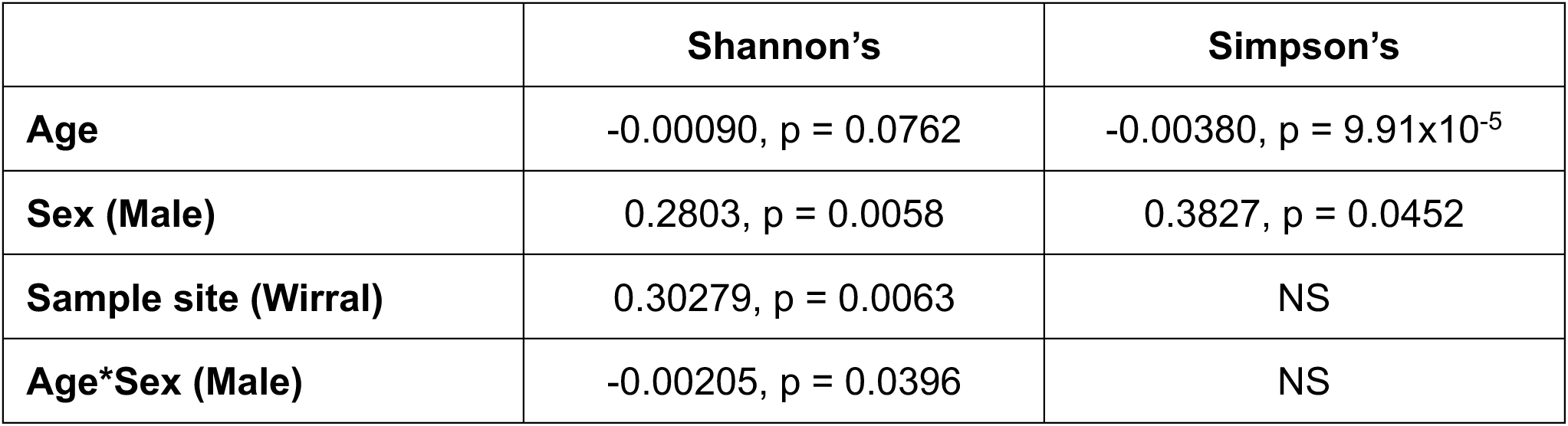
The results of GLM analysis of effects on wild mouse TCR diversity measured using Shannon and Simpson’s diversity indices, of the form VALUE ∼ RECEPTOR TYPE + SITE + AGE + SEX + SEX*AGE, where for each term or the interaction the parameter estimate and the p values are shown. NS is not significant at p = 0.05.

**Supplementary Material 8.**
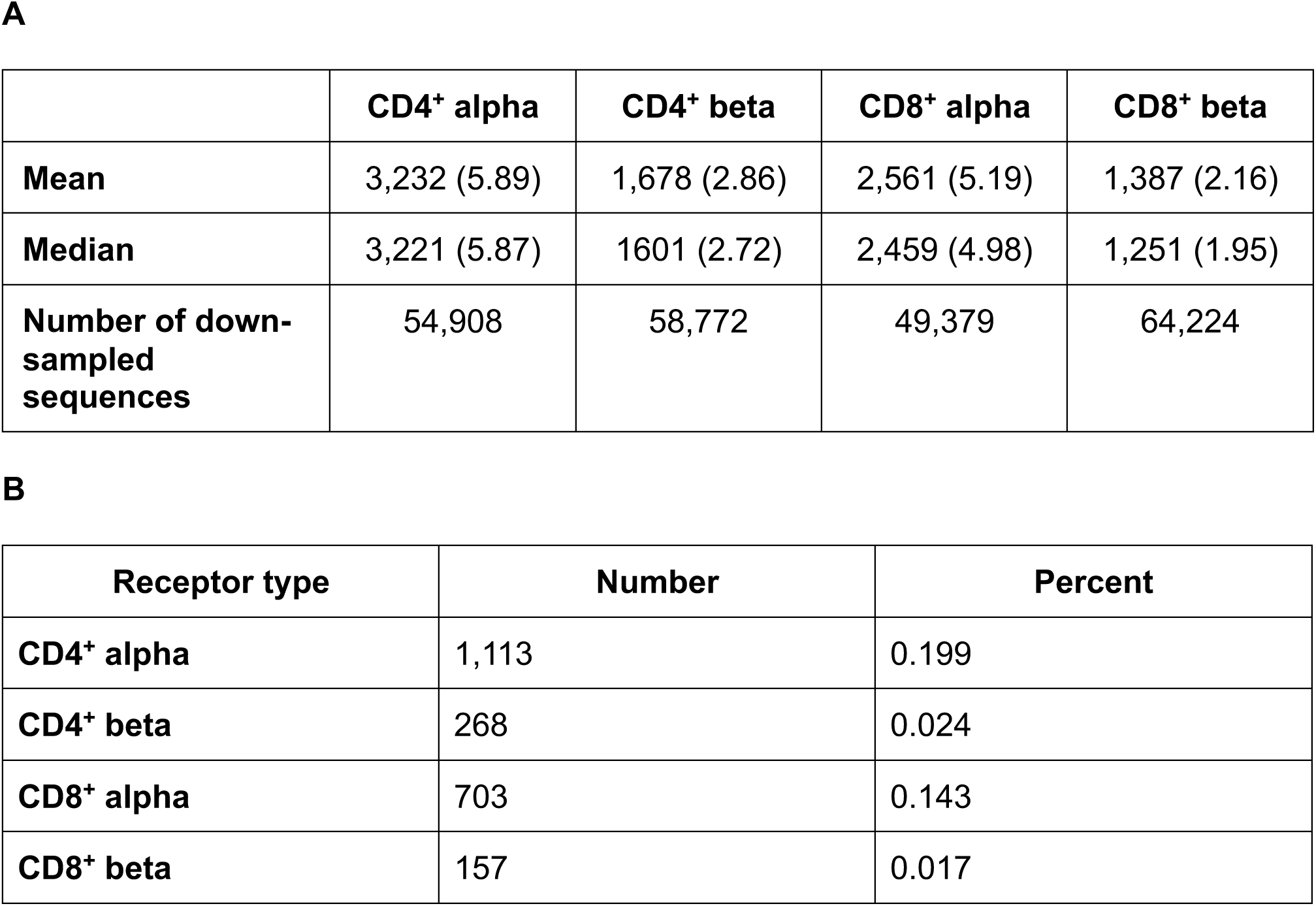
(A) The mean and median number of amino acid-defined TCR sequences shared among wild and laboratory mice and, in parentheses, these expressed as a percentage of the number of down-sampled sequences. (B) The number of amino-acid defined TCR sequences shared among >75 % of mice (both wild and laboratory) and this expressed as a percent of all the unique TCR sequences.

**Supplementary Material 9.**
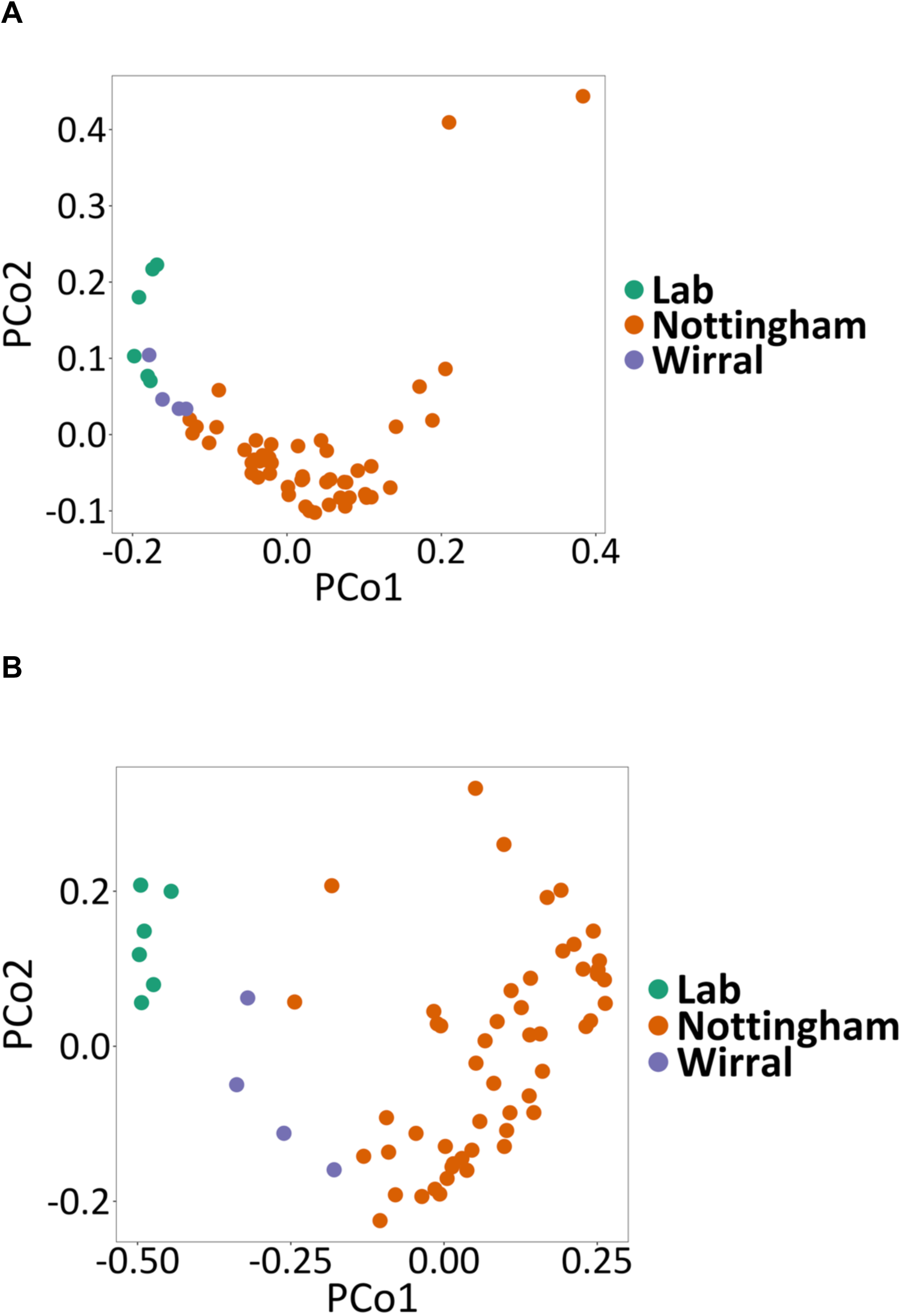
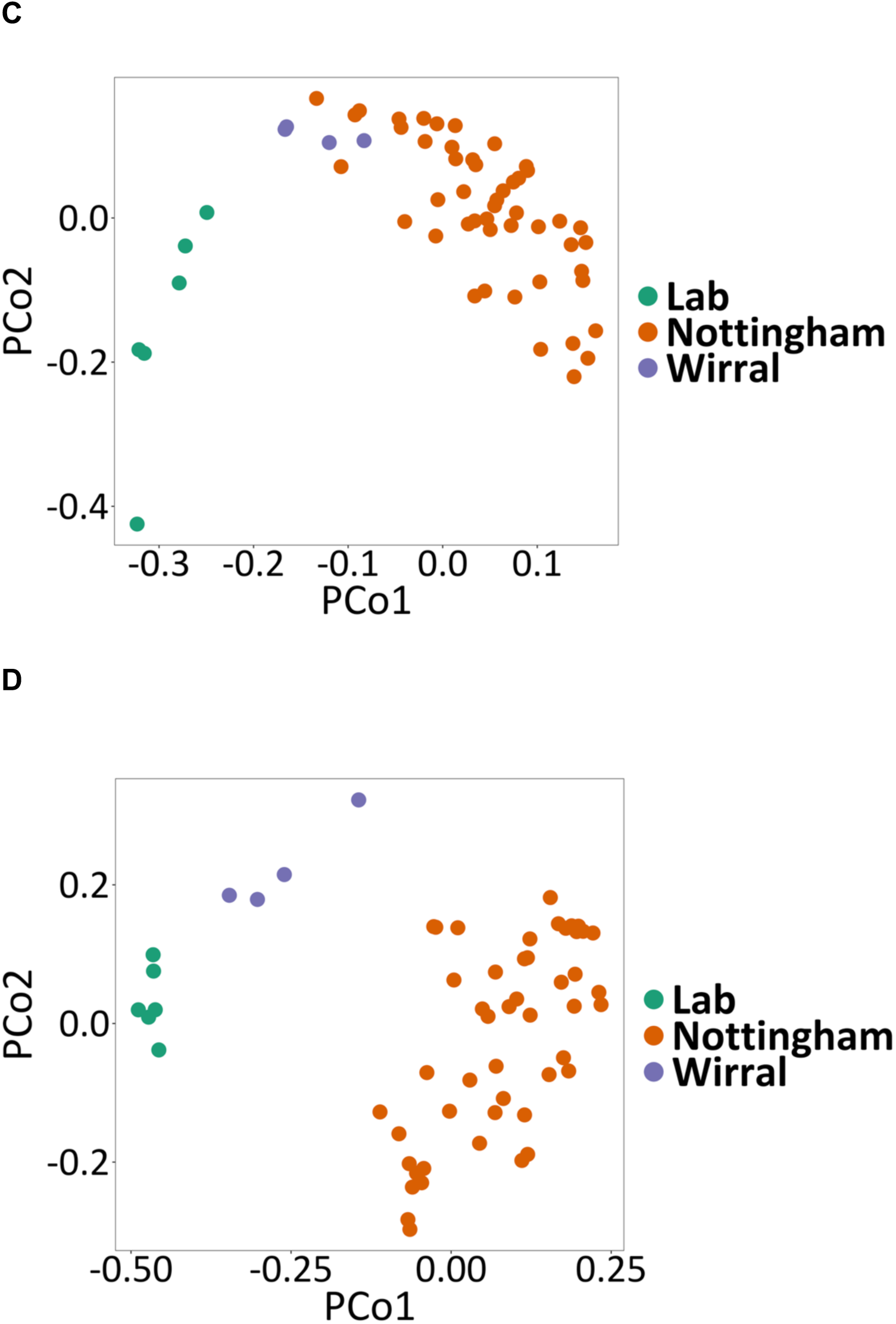
Principal Co-ordinates Analysis (PCoA) showing the first two principal co-ordinates of the number of shared amino acid-defined TCR sequences between all pairwise combinations of wild and laboratory mice, with the origins of the mice colour coded for (A) CD4^+^ alpha, (B) CD4^+^ beta, (C) CD8^+^ alpha and (D) CD8^+^ beta. The percentage of variance explained by PCo1 and PCo2 is for CD4^+^ alpha 11.1 and 9,4; CD4^+^ beta 21.4 and 8.3; CD8^+^ alpha 10.3 and 8.4; CD8^+^ beta 15.5 and 7.5, respectively.

**Supplementary Material 10.**
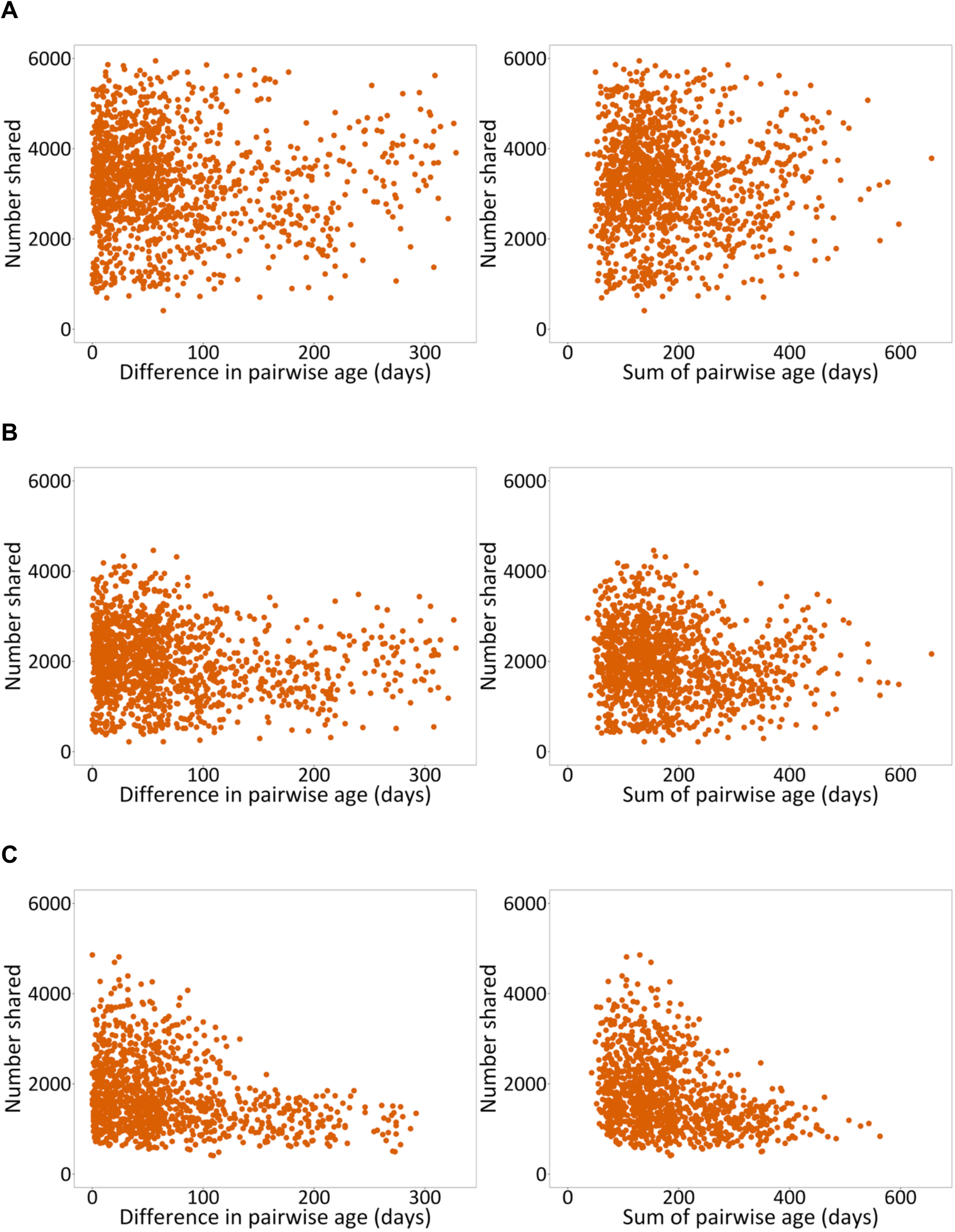
The number of shared amino acid-defined TCR sequences between all pairwise combinations of wild mice from the Nottingham site against the difference in pairwise age (left hand panels) and the sum of the pairwise age (right hand panels), where age is in days for (A) CD4^+^ alpha, (B) CD4^+^ beta and (C) CD8^+^ beta.

**Supplementary Material 11.**
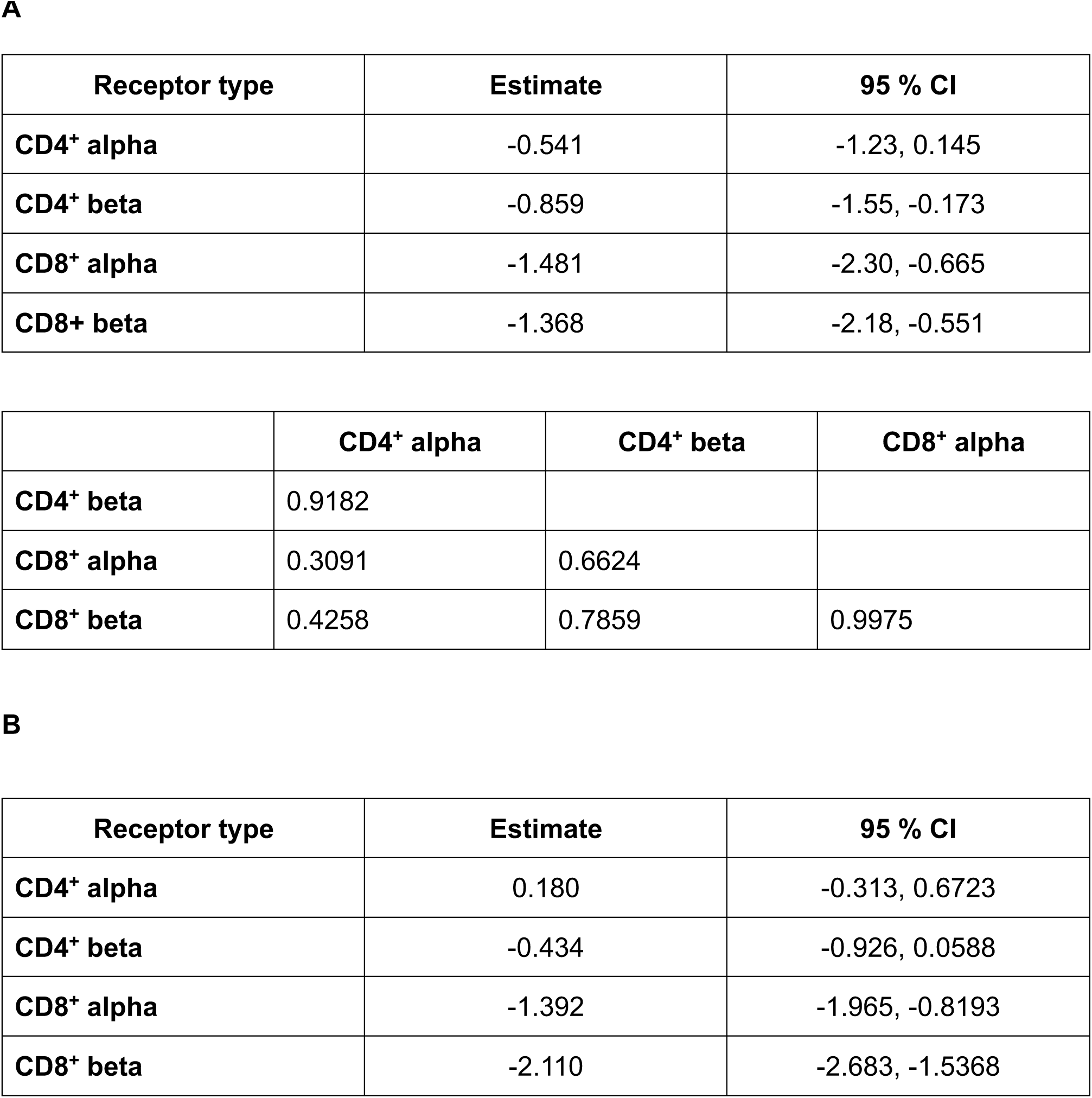
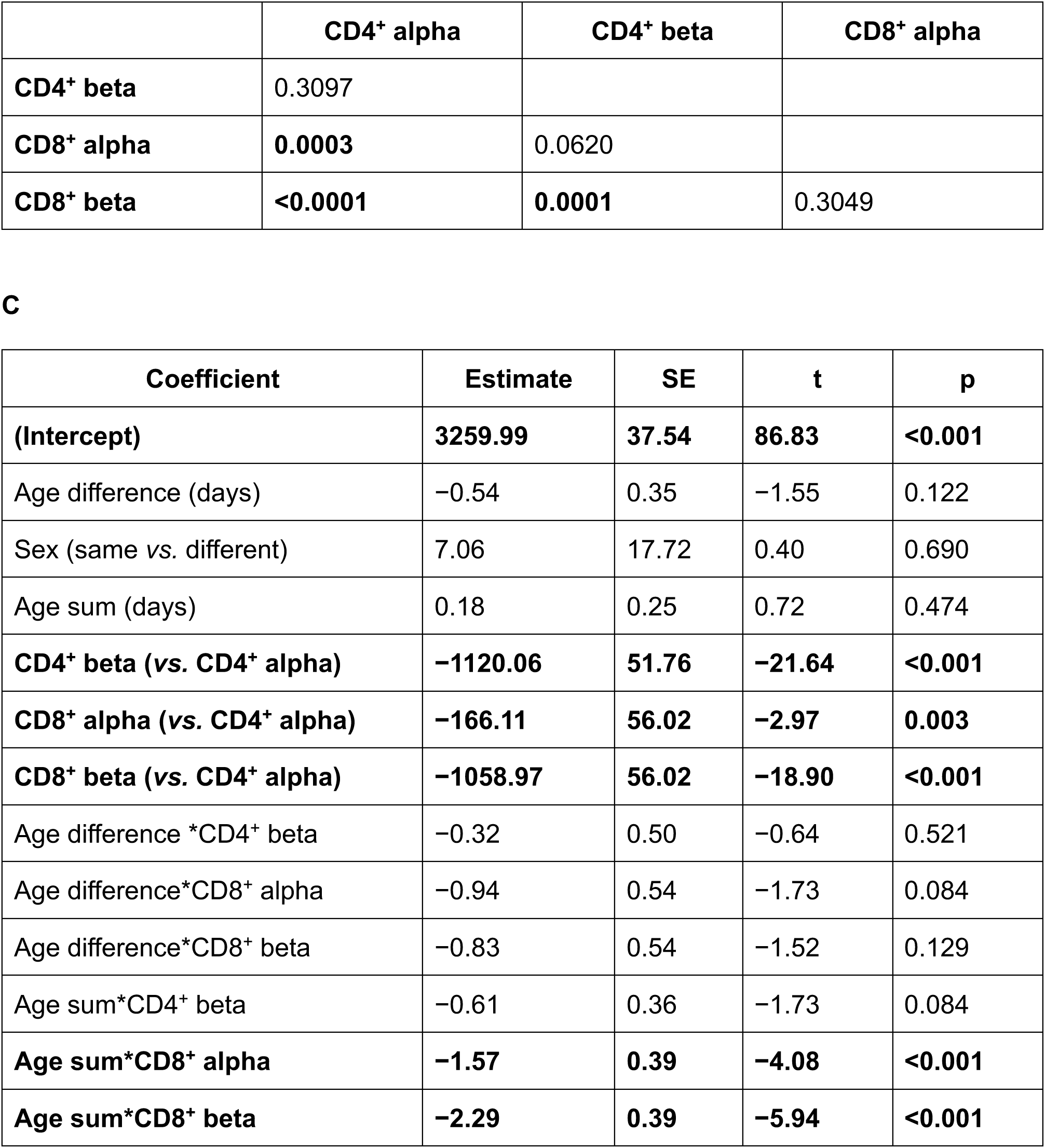
Analysis of the number of shared amino acid-defined TCR sequences and (A) pairwise difference in age and (B) pairwise sum in age of the mice, showing the slope estimates and 95 % confidence intervals (CI) for each receptor type (top table), and the P values for comparisons of the parameter estimates for the four receptor types (bottom table), with bold text showing p < 0.05. (C) Summary of pairwise shared amino acid-defined TCR sequences GLM. Estimates are shown relative to reference levels (CD4^+^ alpha, different sex pairs). Coefficients with p < 0.05 are in bold. GLM details were: *n* = 9,816; residual df = 9,803; null deviance = 1.14e10; residual deviance = 7.55e9; AIC = 160,917.

## Notes

### Competing Interest Statement

The authors have declared no competing interest.

### Summary of Updates

Amendments to ms. following external review.

